# Nuclear compartmentalization of TERT mRNA and TUG1 lncRNA transcripts is driven by intron retention: implications for RNA-directed therapies

**DOI:** 10.1101/2020.07.21.212514

**Authors:** Gabrijela Dumbović, Ulrich Braunschweig, Heera K. Langner, Katarzyna Jastrzebska, Michael Smallegan, Benjamin Blencowe, Thomas R. Cech, Marvin H. Caruthers, John L. Rinn

## Abstract

Numerous global connections have been made between splicing and other layers of gene regulation, including the spatial partitioning of the transcriptome in the cell. Yet, there has been surprisingly little analysis of the spatio-temporal regulation of individual protein-coding and non-coding RNA molecules in single cells. Here we address how intron retention influences the spatio-temporal dynamics of transcripts from two clinically relevant genes: TERT (Telomerase Reverse Transcriptase) pre-mRNA and TUG1 (Taurine-Upregulated Gene 1) lncRNA. Single molecule RNA FISH revealed that nuclear TERT transcripts uniformly and robustly retain two specific introns whose splicing occurs during mitosis. In contrast, TUG1 has a bimodal distribution of fully spliced cytoplasmic and intron-retained nuclear transcripts. We further test the functionality of intron-retention events using RNA-targeting thiomorpholino antisense oligonucleotides to block intron excision. We show that intron retention is the driving force for the nuclear compartmentalization of these RNAs. For both RNAs, altering this splicing-driven subcellular distribution had significant effects on cell growth. Together, these findings show that stable retention of specific introns can orchestrate spatial compartmentalization of RNAs within the cell; this process reveals new targets for RNA-based therapies.

## Introduction

Dynamic regulation of subcellular RNA localization is critical for biological processes ranging from organismal development to cellular activity^1–6^. While the underlying mechanisms have been found for some transcripts, new aspects of RNA localization regulation continue to arise^7,8^. Studies have found that splicing affects RNA localization^9,10^. Over the last years, intron retention has emerged as a regulator of the subcellular distribution and nuclear retention of many messenger RNAs and non-coding RNAs^4,11–16^. Nuclear retention of unspliced or incompletely spliced transcripts can function as a cellular defense mechanism against translation of RNAs with erroneous splicing. However, recent studies show new functions such as buffering protein quantity and rapid response to external stimuli^4,12,13,17^. Some intron retention events can be explained by slow post-transcriptional splicing kinetics^18–23^. Alternatively, very specific introns can be stably retained, adding an additional regulatory layer to RNA functionality^24,25^. For instance, intron retention in long noncoding RNAs (lncRNAs) can give rise to transcripts with unique functions in terms of sequence variability and subcellular distribution^26^. Similarly, some coding RNAs retain specific introns which alters the subcellular localization and availability of their transcripts for translation^27,28^.

Recent technological advances enable spatially resolving and quantifying both coding and non-coding RNA distribution on a single-cell, single-transcript and sub-cellular level^15,29–33^. Here we explored single molecule localization dynamics of two clinically relevant RNAs, TERT mRNA and TUG1 lncRNA, across many human cell types. TERT encodes the catalytic subunit of the ribonucleoprotein complex telomerase, which elongates and maintains telomeres^34^. Telomerase is reactivated in most tumors from almost all cancer types, and it is needed for maintenance of telomeres, which is critical for long-term proliferation of cancer cells^35,36^. The *TERT* gene is silenced in differentiated cells, hence TERT has been considered as a promising therapeutic target in cancer^37,38^. We and others recently showed that the majority of TERT transcripts are compartmentalized in the nucleus^39,40^, suggesting a potential regulatory mechanism acting at the level of RNA. TUG1 lncRNA has a role in many cellular processes and is associated with malignancies, where it has an oncogenic role (inferred as onco-lncRNA)^41–49^. We and others showed that TUG1 lncRNA is located in both the nucleus and cytoplasm and, correspondingly, it was shown to have a function in both compartments^31,50,51^. Together, TERT and TUG1 transcripts provide models to elucidate when and where splicing occurs, which introns are retained and how this in turn affects the cellular state.

We address these questions using single molecule RNA FISH (smRNA FISH) combined with modified antisense oligonucleotides (ASOs) that localize to the nucleus and direct specific splicing events. We observe that retention of specific introns drives the nuclear retention of both TERT pre-mRNA and TUG1 lncRNA transcripts. In the case of TERT transcripts, we find that intron retention is regulated during the cell cycle, with two specific introns retained during interphase and spliced out during mitosis. For TUG1 transcripts, we observed two distinct populations of fully spliced, cytoplasmic and intron-retained, nuclear RNAs. Our results further show that nuclear TUG1 and TERT transcripts are more stable than the corresponding cytoplasmic transcripts. We tested the functional significance of these intron-retention events using ASOs that further drive intron retention and result in a clear shift in subcellular localization. Altering the nuclear-cytoplasmic distribution of TERT and TUG1 transcripts had significant functional consequences on a cellular scale and reduced cell growth *in vitro*. Collectively, our findings provide new evidence for the importance of spatio-temporal regulation of intron retention and suggest a novel approach to intervene in RNA-based therapies with modified antisense oligos.

## Results

### 1. Nuclear TUG1 lncRNA and TERT mRNA retain introns

We and others previously observed that TERT transcripts are unexpectedly more abundant in the nucleus than the cytoplasm^39,40^. Somewhat similarly, smRNA FISH revealed that the TUG1 lncRNA is evenly distributed between the nucleus and cytoplasm (Fig. 1)^31,51^. We sought to determine molecular features or splicing patterns that could differentiate nuclear versus cytoplasmic localization of these transcripts. By analyzing available RNA-Seq data from the ENCODE consortium^52^, high read coverage across both TUG1 introns was observed (Fig. 1a and Extended data Fig. 1a), while TERT had high read coverage across two of its introns, intron 11 and intron 14 (Fig. 1b and Extended data Fig. 1b). Next, we calculated the splicing efficiency of each intron using published RNA-Seq data from human induced pluripotent stem (iPS) cells. Intron-retention events were calculated as percent intron retention (PIR) using vast-tools^53^ as described previously^11^. Briefly, intron retention was evaluated as the ratio of read counts mapping to exon-intron junctions relative to the total number of exon-intron junction reads plus spliced exon-exon junction reads (see Methods). The results show that TERT specifically retains introns 11 and 14 in iPS cells (PIR of 30% and 31%, respectively) while the other introns are efficiently spliced (Fig. 1c and Extended data Table 1). Further, we confirmed the retention of the first intron in TUG1; it has a PIR of 46% in iPS cells whereas TUG1 intron 2 is absent from the VastDB human database.

**Figure 1:**
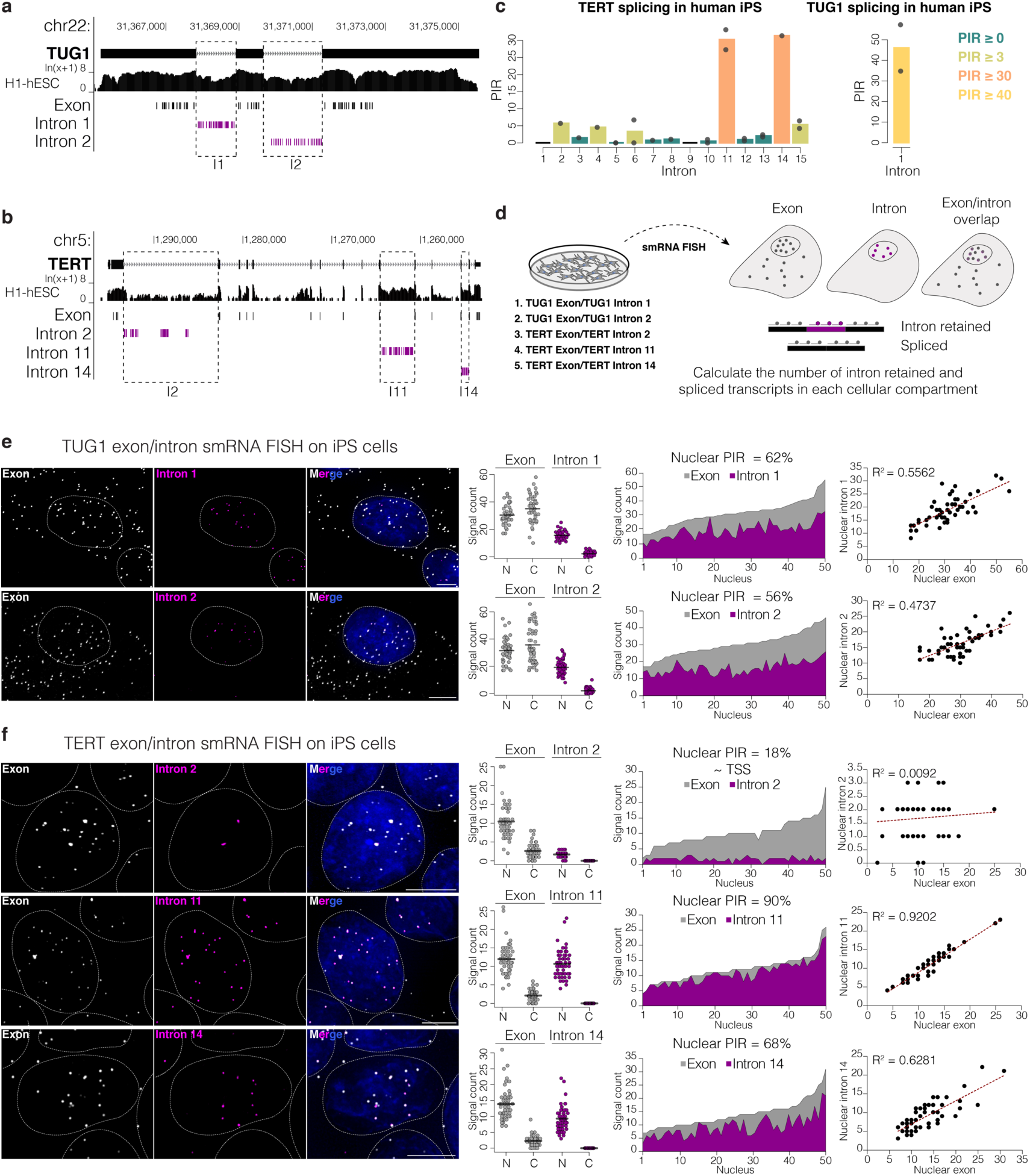
Retention of specific introns correlates with nuclear localization of TERT mRNA and TUG1 lncRNA in hES/iPS cells. **a**, UCSC Genome Browser showing the TUG1 locus (hg19) and the RNA-seq track from human ES cells from ENCODE. Below, the location of probes used in smRNA FISH. Exon probes, grey; intron probes, magenta. **b**, UCSC Genome Browser showing the TERT locus (hg19) and the RNA-seq track from human ES cells from ENCODE. Below, the location of probes used in smRNA FISH. Exon probes, grey; intron probes, magenta. **c**, Percentage of intron retention of TERT (left) and TUG1 (right) in human iPS cells obtained with vast-tools analysis of RNA-seq data. Bars, means across replicates; dots, individual replicates. Introns with insufficient read coverage are shown as black lines in TERT plot. TUG1 intron 2 was absent in the VastDB database. **d**, SmRNA FISH scheme. Co-localizing exon and intron signals are considered as unspliced, exon-only signal as spliced. **e**, Maximum intensity projections of representative images of TUG1 exon/intron smRNA FISH on iPS cells. Exon, gray; intron 1 and 2, magenta. Nucleus, blue, outlined with a dashed circle. Scale bar, 5 μm. Towards the right: quantification (*n* = 50) of spliced and unspliced transcripts for each intron in the nucleus (N) and cytoplasm (C); average percentage of nuclear intron retention (nuclear PIR) of each intron; correlation between nuclear intron and nuclear TUG1 quantity. **f**, Maximum intensity projections of representative images of TERT exon/intron smRNA FISH on iPS cells. Exon, gray; introns 2, 11 and 14, magenta. Nucleus, blue, outlined with a dashed circle. Scale bar, 5 μm. Towards the right: quantification (*n* = 50) of spliced and unspliced transcripts for each intron; average nuclear PIR of each intron; correlation between nuclear intron and nuclear TERT quantity.

We hypothesized that the transcripts with retained introns would be nuclear localized. To determine RNA localization, we designed smRNA FISH probes tilling across TUG1 and TERT exons and introns (Fig. 1a and Fig. 1b). Dual-color smRNA FISH probes independently targeted TUG1 exons and intron 1 or intron 2, and TERT exons and intron 11 or intron 14 (Fig. 1d). We further applied smRNA FISH against TERT intron 2 and GAPDH intron 2 as controls for non-retained introns. Imaging datasets were processed, and co-localized exon and intron spots were quantified as intron-retaining transcripts, while exon-only signal was quantified as transcripts with that specific intron spliced out (Fig. 1d).

RNA imaging on a human iPS cell line showed an even distribution of TUG1 in the nucleus and cytoplasm (average ∼48% and ∼52%, respectively) (Fig. 1e). The vast majority of the transcripts in the nuclear fraction had retained introns, whereas those in the cytoplasmic fraction did not (Fig. 1e). More specifically, the average percentage of nuclear intron retention (nuclear PIR, expressed as percentage of nuclear intron-retaining transcripts over total nuclear transcripts) for TUG1 intron 1 and intron 2 in iPS cells was 62% and 56%, respectively, with significant correlations between the magnitude of detected intron retention and nuclear TUG1 transcript levels (R^2^=0.56 intron 1, R^2^=0.47 intron 2; *P*=5.16 × 10^−10^ and *P*=3.33 × 10^−8^, respectively, Pearson correlation, Fig. 1e).

TERT transcripts are retained in the nucleus to an even higher degree than TUG1 transcripts, with on average 86% of total detected TERT RNAs retained in the nucleus of iPS cells (Fig. 1f). Intron 11 of TERT has a high nuclear PIR (90%) which correlates with the quantity of detected nuclear TERT RNA (R^2^=0.92, *P*< 2.2 × 10^−16^, Pearson correlation). Intron 14 was also retained, albeit at a lower proportion (nuclear PIR=68%), and it also showed a significant correlation with the quantity of nuclear TERT (R^2^=0.63, *P*=7.1 x-10^−12^, Pearson correlation). These results indicate that TERT intron 11 might have a greater impact on the nuclear retention of TERT RNA than intron 14. TERT intron 2, a control for a non-retained intron in iPS cells, had average nuclear PIR=18% with no correlation with the quantity of nuclear TERT (R^2^=0.0092, *P*=0.48, Pearson correlation). GAPDH intron 2 smRNA FISH showed on average of between 1 and 2 punctate signals per cell, which overlapped with GAPDH exon signal and marked the active transcription sites, hence further supporting the specificity of the smRNA FISH approach detecting intron retention in TUG1 and TERT (Extended data Fig. 1c).

### 2. TUG1 and TERT intron retention across cancer cell types

The pattern of nuclear localization and intron retention observed in healthy iPS cells that endogenously express TUG1 and TERT led us to explore whether this phenomenon is specific to iPS cells or also occurs in other cell types and contexts such as cancer, where TERT expression is reactivated and TUG1 is expressed. We performed the same analysis for TUG1 on four cancer cell lines (osteosarcoma U-2 OS, cervical cancer HeLa, colorectal cancer HCT116, and glioblastoma LN-18) and two non-tumor-derived cell types (embryonic kidney HEK293T and BJ fibroblasts); and for TERT on 4 cell lines with TERT re-activation (HeLa, HCT116, HEK293T and LN-18).

RNA-Seq analysis showed high read coverage across TUG1 intron 1 and 2 (Fig. 2a) in all cell lines, indicating that a large fraction of this lncRNA has retained introns. Our smRNA FISH revealed a consistent nuclear/cytoplasmic localization for TUG1 regardless of the cell or cancer type (Fig. 2b and Extended data Fig. 2a for fibroblasts). As before, there was a significant correlation between the quantity of nuclear TUG1 and intron retention (R^2^ ≥0.5, *P*<0.001 in all cell lines tested, Pearson correlation) (Fig. 2b and Extended data Fig. 2a for fibroblasts). There were modest differences in the nuclear PIR between cell lines for both intron 1 and intron 2 (mean value ranging from 52% to 75% for intron 1, and from 52% to 67% for intron 2) (Fig. 2c left). To determine whether the retention of the introns was correlated with the overall nuclear retention of TUG1, we compared total PIR with nuclear enrichment of TUG1 (expressed as percentage of nuclear TUG1 RNA over total TUG1 per cell). Cell lines with higher total PIR of intron 1 and intron 2 tended to have more nuclear TUG1, indicating a correlation between the extent of TUG1 intron retention and nuclear localization (R^2^=0.86 for nuclear enrichment vs. total PIR intron 1; R^2^=0.93 for nuclear enrichment vs. total PIR intron 1, *P*=0.0015 and *P*=0.0004, respectively, Pearson correlation) (Fig. 2c right and Extended data Fig. 2b). Overall, TUG1 showed dual localization and retention of both introns across all analyzed cell lines, with corresponding differences in the ratios of nuclear vs. cytoplasmic TUG1 transcripts and PIR values between cell lines, thereby opening the possibility that this process is being fine-tuned and regulated in a cell type dependent manner.

**Figure 2:**
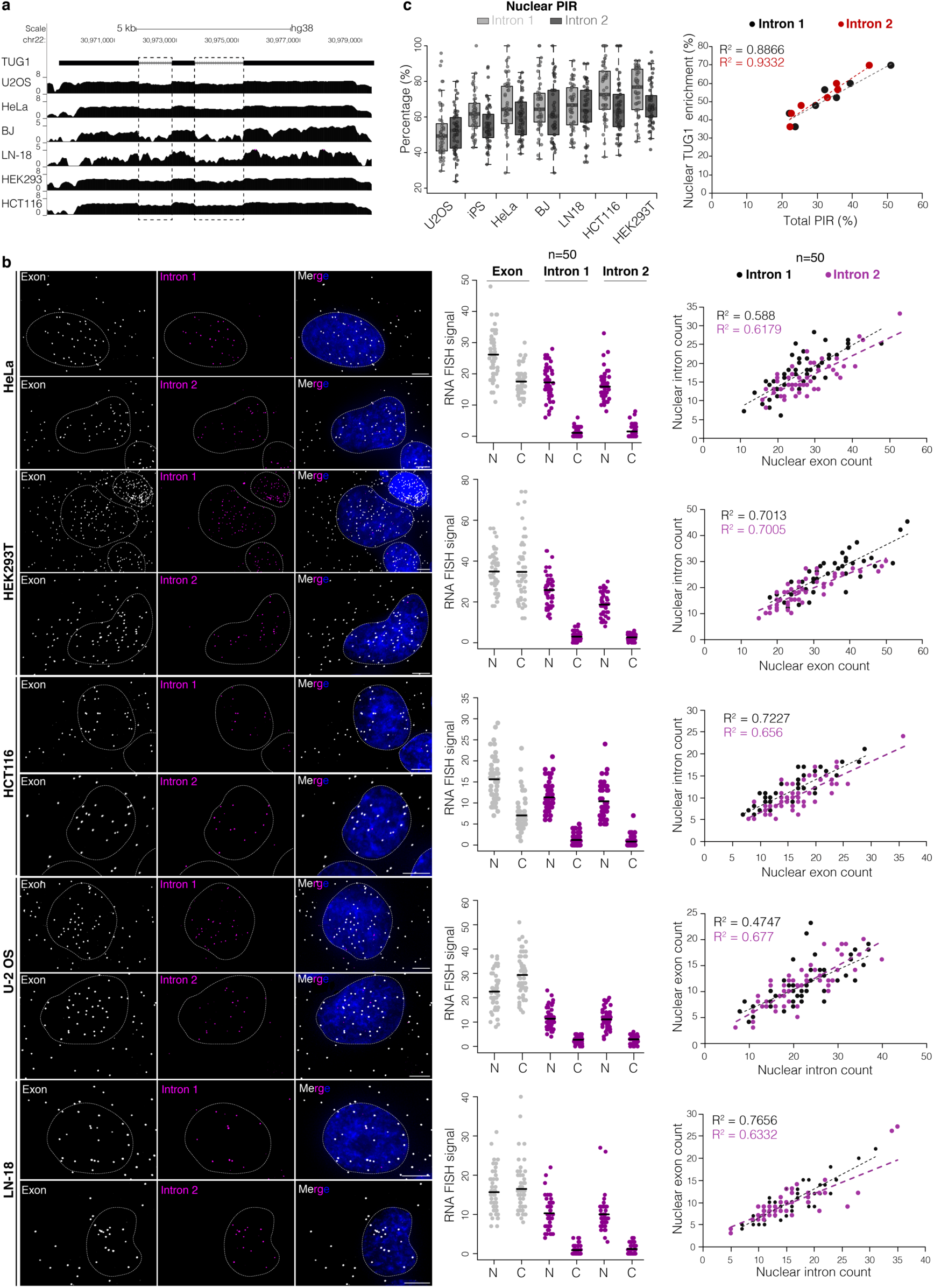
TUG1 intron retention is common and fluctuates across cells lines. **a**, UCSC Genome Browser showing RNA-seq coverage across TUG1 locus (hg38) from multiple cell lines. Scale ln(x+1). **b**, Maximum intensity projections of representative images of TUG1 exon/intron smRNA FISH across different cell lines. Exon, gray; intron 1, 2, magenta; nucleus, blue, outlined with a dashed line. Scale bar, 5 μm. Middle: quantification (*n* = 50) of spliced and unspliced transcripts for each intron in the nucleus (N) and cytoplasm (C). Right: correlation between nuclear intron and nuclear TUG1 quantity, intron 1, black; intron 2, magenta. **c**, Nuclear PIR for each intron across cell lines. Right: correlation between TUG1 nuclear enrichment and total PIR between different cell lines. Each data point, mean value from one cell line, all measurements shown in Extended data Fig. 2b. Intron 1, black; intron 2 red.

We next explored TERT intron retention across cell lines in a similar manner as for TUG1 (Fig. 3). RNA-Seq analysis showed high read coverage in introns 11 and 14 (and in HCT116 cells, intron 2), suggesting their retention (Fig. 3a). Turning to smRNA FISH, the HCT116, HEK293T and LN-18 cell lines expressed TERT in the majority of cells (Fig. 3b). The LN-18 and HEK293T cell lines showed similar nuclear enrichment of TERT as the iPS cell line (average, 82% and 89% TERT RNA in the nucleus, respectively) (Fig. 3c). This was accompanied by high retention of intron 11 (nuclear PIR=91% and 89% for LN-18 and HEK293T, respectively). Retention of intron 11 had a significant correlation with the quantity of nuclear TERT (R^2^=0.94 and 0.96 for LN-18 and HEK293T, respectively, *P*< 2.2 × 10^−16^ for both cell lines, Pearson correlation). As in iPS cells, intron 14 was retained to a lesser extent (nuclear PIR=61% and 47% for LN-18 and HEK293T, respectively) with a modest, but significant, correlation with the quantity of nuclear TERT (R^2^=0.28 and 0.67 for LN-18 and HEK293T, respectively, *P*=7.0 × 10^−5^ and 2.8 × 10^−13^, respectively, Pearson correlation). HeLa cells were excluded from this analysis because they had very few detectable molecules of TERT RNA per cell (Extended data Fig. 3).

**Figure 3:**
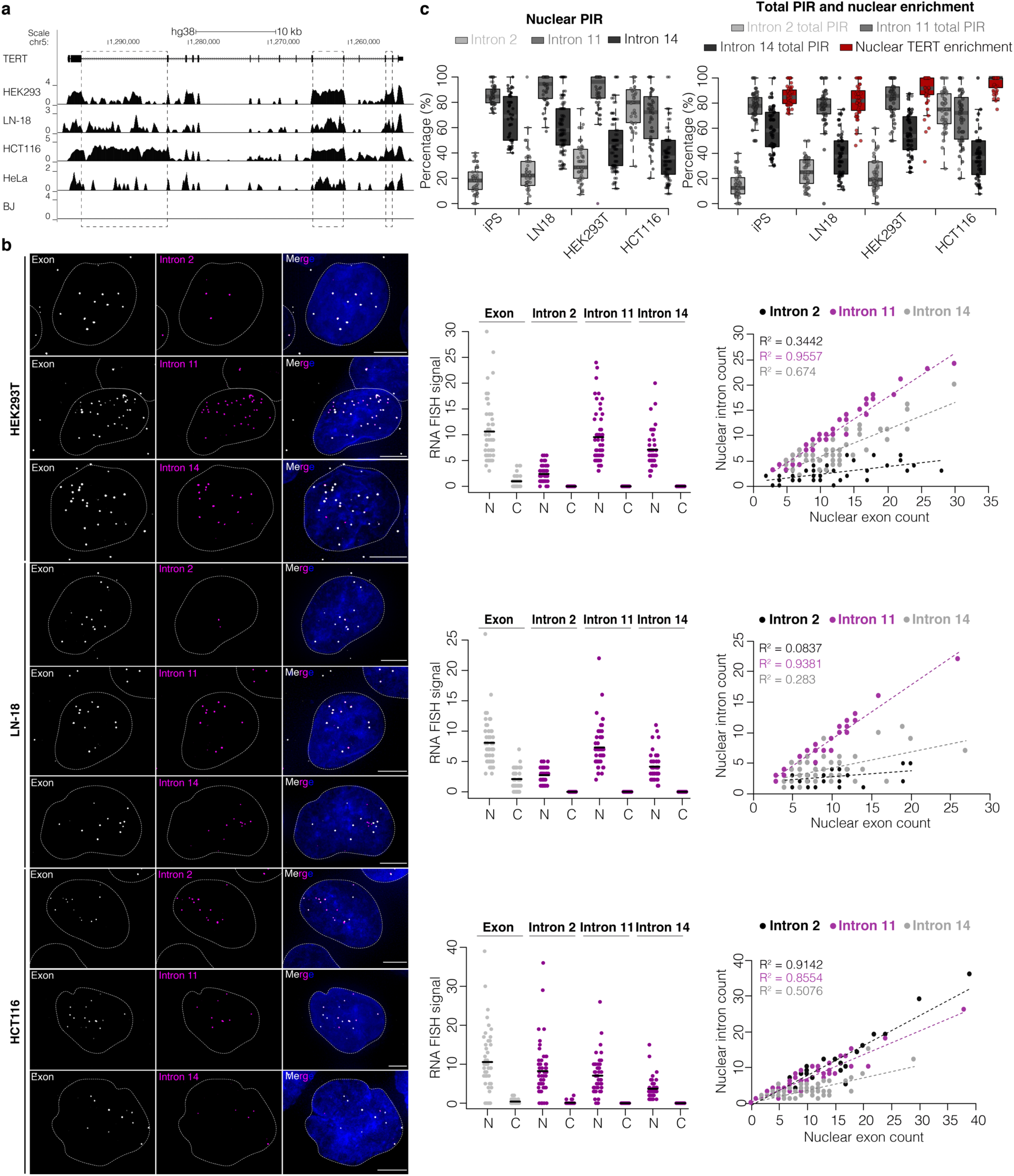
Retention of TERT intron 11 is robust across cell lines. **a**, UCSC Genome Browser showing RNA-seq coverage across TERT locus (hg38) from multiple cell lines. Scale ln(x+1). **b**, Maximum intensity projections of representative images of TERT exon/intron smRNA FISH across different cell lines. Exon, gray; introns 2, 11, 14, magenta; nucleus, blue, outlined with a dashed line. Scale bar, 5 μm. Middle: quantification (*n* = 50) of spliced and unspliced transcripts for each intron in the nucleus (N) and cytoplasm (C). Right: correlation between nuclear intron and nuclear TERT count; intron 2, black; intron 11, magenta; intron 14, gray. **c**, Nuclear PIR for each intron across cell lines. On the right: total PIR of each intron and percentage of nuclear enrichment of TERT across cell lines.

The HCT116 cell line showed some differences compared to iPS, LN-18 and HEK293T cell lines. First, very rarely were spliced TERT transcripts detected in the cytoplasm; on average 96% were found in the nucleus (Fig. 3b and 3c). While intron 2 was not significantly retained in other cell lines examined, HCT116 retained intron 2 in nuclear TERT (nuclear PIR=78%, R^2^=0.91 with quantity of nuclear TERT, *P*<2.2 × 10^−16^, Pearson correlation), alongside intron 11 (nuclear PIR=69%, R^2^=0.86 with quantity of nuclear TERT, *P*<2.2 × 10^−16^, Pearson correlation), while intron 14 was retained less efficiently (nuclear PIR=40%, R^2^=0.50 with quantity of nuclear TERT, *P*=6.5 × 10^−9^, Pearson correlation). This atypical retention of intron 2 can also be observed in the corresponding RNA-seq for HCT116 (Fig. 3a).

Based on our analysis, we find intron 11 robustly retained across different cell lines, while intron 14 showed less and more variable retention, similar to what was observed in iPS cells. Furthermore, these data illustrate the need to analyze possible splicing aberrations that might influence subcellular localization of TERT in cancer, as shown here in HCT116 cell line.

### 3. TUG1 and TERT intron retention is conserved across species

We reasoned that if these specific intron retention events for TUG1 and TERT were biologically relevant, they would show evolutionary conservation. To address this, we performed several analyses between human and mouse. The TUG1 locus has high sequence conservation between human and mouse, maintaining the same gene organization (3 exons and 2 introns) and exhibiting 62.5% overall sequence conservation, 70.2% in exons and 52.2% in introns (Extended data Fig. 4a). We observed that the 5’ exon has slight differences in the annotation compared to human TUG1 (smaller than the corresponding exon in human) (Extended data Fig. 4b). Similarly, the TERT locus maintains the same gene organization (16 exons and 15 introns) between human and mouse (Extended data Fig. 4a). The overall nucleotide sequence conservation between human and mouse TERT loci is only 29.1%, which is mostly due to the low intron sequence conservation (25.5%) while coding sequences exhibit higher conservation (60.7%). Thus, TUG1 and TERT show similar evolutionary conservation in their exonic structures.

**Figure 4:**
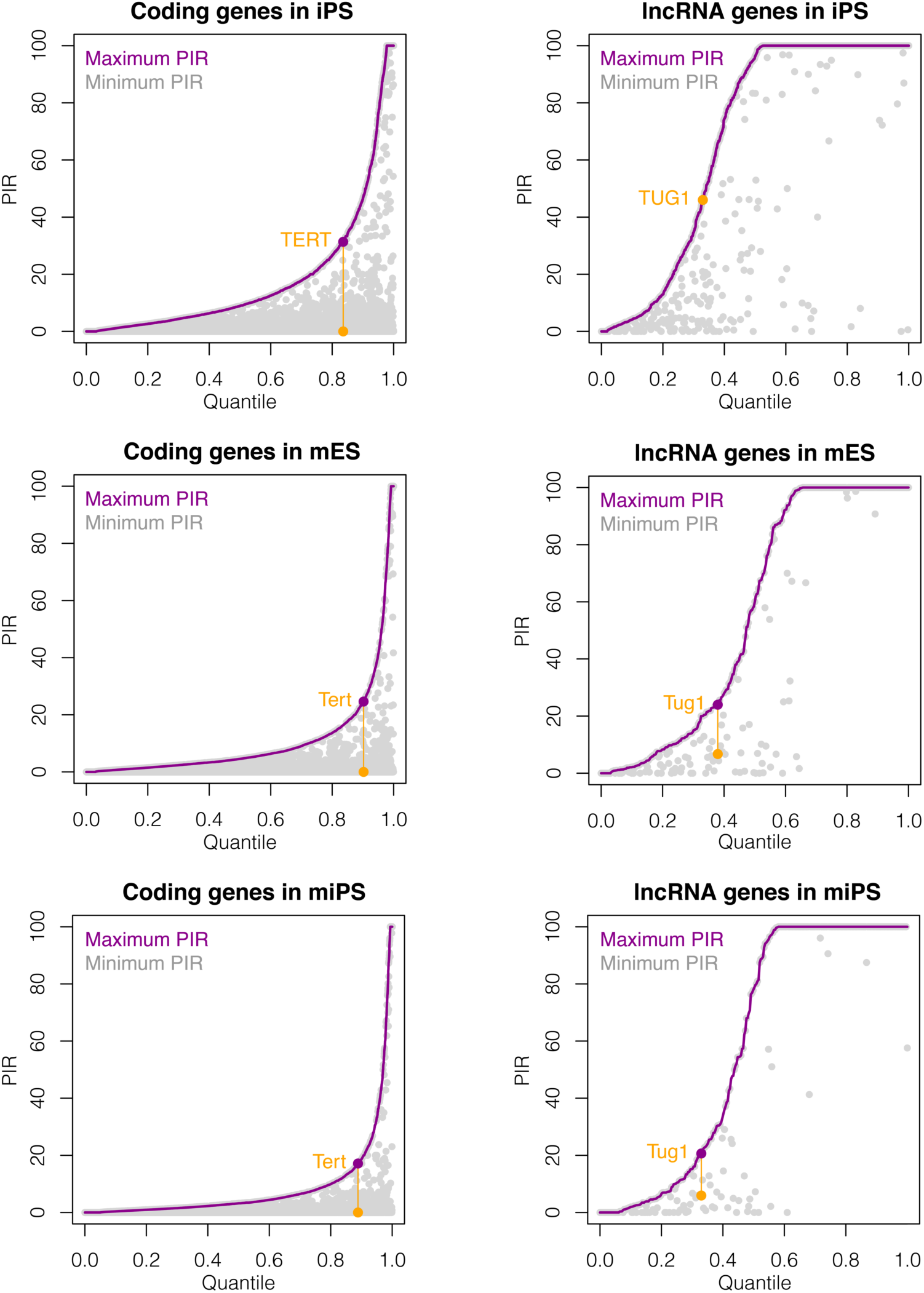
Global PIR analysis for coding and lncRNA genes. Cumulative distribution of maximum PIR levels for each coding and lncRNA gene in hiPS, mES and miPS cells (in purple). Minimum PIR value for the same gene is plotted in grey at the same x-axis position. Introns with maximum and minimum PIR values from TERT and TUG1 are connected with a yellow line.

We sought to determine whether intron retention is a conserved phenomenon in TUG1 and TERT transcripts in comparable cell types across species. We analyzed the splicing efficiency of individual Tert and Tug1 introns using published RNA-Seq data from mouse iPS (miPS) and mouse embryonic stem (mES) cells (Extended data Fig. 4b). In mES and miPS cells, Tug1 intron 1 is not highly retained (PIR of ∼6% in miPS and mES), while mouse intron 2 had a higher PIR of ∼20%. We applied smRNA FISH to further determine retention of both introns and subcellular localization patterns of Tug1 in mES cells (Extended data Fig. 4c). We observed conserved dual localization of Tug1 in the nucleus and cytoplasm (average 61% nuclear Tug1). Intron 2 was highly retained in nuclear Tug1 (PIR=62%). However, intron 1 is less retained (PIR=24%), thereby validating the more efficient splicing of intron 1 in mouse compared to human TUG1.

Splicing efficiency analysis of Tert introns showed efficient splicing of intron 11 and 14 in mouse (PIR=3.4% and 0%, respectively), contrary to their high retention in human cells (PIR=30.2% and 31.4%, respectively) (Extended data Fig. 4b). In contrast, intron 3 and intron 7 were highly retained in mouse Tert (PIR=24.6% and 23%, respectively, in mES, and 13.3% and 17.2%, respectively, in miPS).

We next analyzed intron features that could potentially discriminate retained from efficiently spliced introns. Previously, it was shown that retained introns are significantly associated with elevated CG content, reduced length, and relatively weak donor and acceptor splice sites^28^. Introns 3, 7, 11 and 14 are in general longer in human than mouse (Extended data Table 2). No significant differences in GC content were found, except for intron 7 having lower GC content in mouse. We further analyzed the conservation and strength of splice sites of all TERT introns, focusing on highly retained TERT introns in either human or mouse (intron 3, 7, 11 and 14). In all instances, the canonical GT-AG pair is present (Extended data Fig. 5). The acceptor and donor splice sites are classified as strong, with no significant differences in the strength of splice sites correlating with intron retention, with the exception of intron 7 which has a weaker donor splice site in mouse. Extensive deletions and SNPs +2 bp downstream of donor splice sites and upstream of the acceptor sites are present, which opens the possibility of binding a different plethora of RNA binding proteins between human and mouse. Considering the strength of donor and acceptor splice sites and highly efficient splicing of retained human TERT introns during mitosis (shown below), it seems probable that during interphase the excision of those introns is prevented by binding of splicing repressors, which is relieved in mitosis. In contrast, TUG1 introns are highly conserved between human and mouse in length, GC content and splice site strength (Extended data Fig. 5, Extended data Table 2).

**Figure 5:**
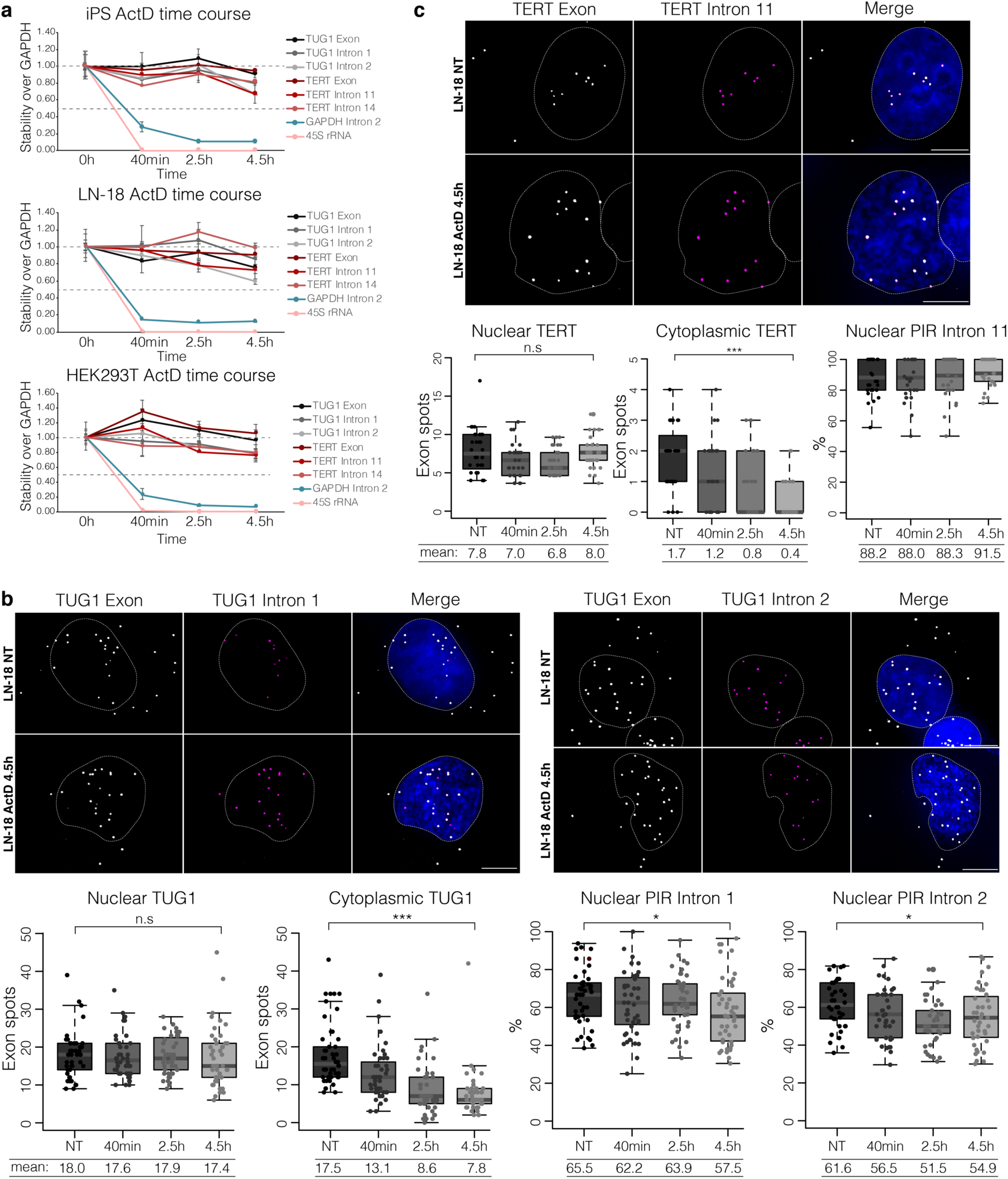
Intron-retained nuclear TUG1 and TERT are long-lived transcripts, stably retained in the nucleus. **a**, Relative stability of TUG1 and TERT exons and introns compared to GAPDH mRNA measured by RT-qPCR in iPS, HEK293T and LN-18 cells during a 4.5 h ActD time course. GAPDH intron 2, a control for an efficiently spliced intron; 45S rRNA, a control for a precursor RNA. **b**, Maximum intensity projection of LN-18 smRNA FISH targeting TUG1 exon (gray) and intron 1 (magenta) or intron 2 (magenta) at time point 0 (NT) and 4.5 h after ActD treatment. Scale bar, 5 μm. Below, smRNA FISH quantification (*n* = 30-50) at each time point of spliced and unspliced TUG1 transcripts in the nucleus and cytoplasm; PIR of intron 1 and intron 2 at each time point. n.s. = not significant, \**P* ≤ 0.05, \*\*\**P* ≤ 0.001, as evaluated by unpaired *t*-test versus NT. **c**, Maximum intensity projection of LN-18 smRNA FISH targeting TERT exon (gray) and intron 11 (magenta) at time point 0 (NT) and 4.5 h after ActD treatment. Scale bar, 5 μm. Below, smRNA FISH quantification (*n* = 30-50) at each time point of spliced and unspliced TERT transcripts in the nucleus and cytoplasm; nuclear PIR of intron 11 at each time point. n.s. = not significant, \*\*\**P* ≤ 0.001, as evaluated by unpaired *t*-test versus NT.

Next we sought to determine whether the observed retention of specific introns in TERT and TUG1 is atypical or a common phenomenon among coding and lncRNA genes. We analyzed PIR of each intron for every mRNA and lncRNA across hiPS, mES and miPS cells (Fig. 4 and Extended data Table 3,4). For each gene, we plotted the maximum PIR among all introns in a given transcript, together with the minimum PIR, if applicable. Intron retention is generally high in lncRNA genes, extending previous observations that introns in UTRs and non-coding genes were particularly highly retained^11^. Interestingly, many coding genes have at least one retained intron as well as fully spliced introns. In both human and mouse, TUG1 appears to have a maximum PIR typical for a lncRNA gene. On the other hand, PIR of TERT retained introns is within the top 20% of coding genes for both species.

Collectively, these results indicate that the TERT and TUG1 intron retention phenomenon is conserved across species; where in case of TERT it is not tied to specific introns, which strengthens the possibility that intron retention is relevant for TERT regulation.

### 4. Intron-retained nuclear TUG1 and TERT are stable transcripts that remain in the nucleus after transcription inhibition

Some RNA intermediates retain certain introns due to slow post-transcriptional splicing kinetics^12,23^. To test whether TUG1 and TERT retain introns due to slow splicing kinetics or if those are stable transcripts, we treated cells with Actinomycin D (ActD). ActD inhibits Pol I, Pol II and Pol III by intercalating in the DNA and preventing transcription elongation^54,55^. Cell lines that endogenously co-express TUG1 and TERT (iPS, LN-18, HEK293T) were treated for up to 4.5 hours and harvested for RT-qPCR at several time points (0 h, 40 min, 2.5 h and 4.5 h). We monitored the stability of TUG1 and TERT exons and retained introns. As a stable RNA control, we used GAPDH, while GAPDH intron 2 and pre-ribosomal RNA (45S rRNA) were used as controls for nascent RNAs. We observed an immediate decrease in the nascent 45S rRNA and GAPDH intron 2 after 40 min of ActD treatment. In contrast, in healthy and cancer cell lines, intron-containing TUG1 and TERT RNAs were highly stable even after 4.5 h of transcription inhibition (Fig. 5a).

We used smRNA FISH during ActD treatment to determine the stability and spatial localization of intron-retained and spliced TUG1 and TERT. Specifically, nuclear TUG1 remained stable across the ActD time course (Fig. 5b for LN-18 and Extended data Fig. 6a for iPS). In contrast, cytoplasmic TUG1 gradually decreased in both cell lines during ActD time points (∼2.2-fold decrease, *P*≤0.001, unpaired *t*-test for LN-18; ∼2-fold decrease, *P*≤0.001, unpaired *t*-test for iPS). Furthermore, retention of intron 1 and 2 remained high even after 4.5 h of treatment, and unspliced TUG1 remained nuclear. Thus, the nuclear, intron-retained TUG1 fraction is more stable than the fully spliced cytoplasmic fraction.

**Figure 6:**
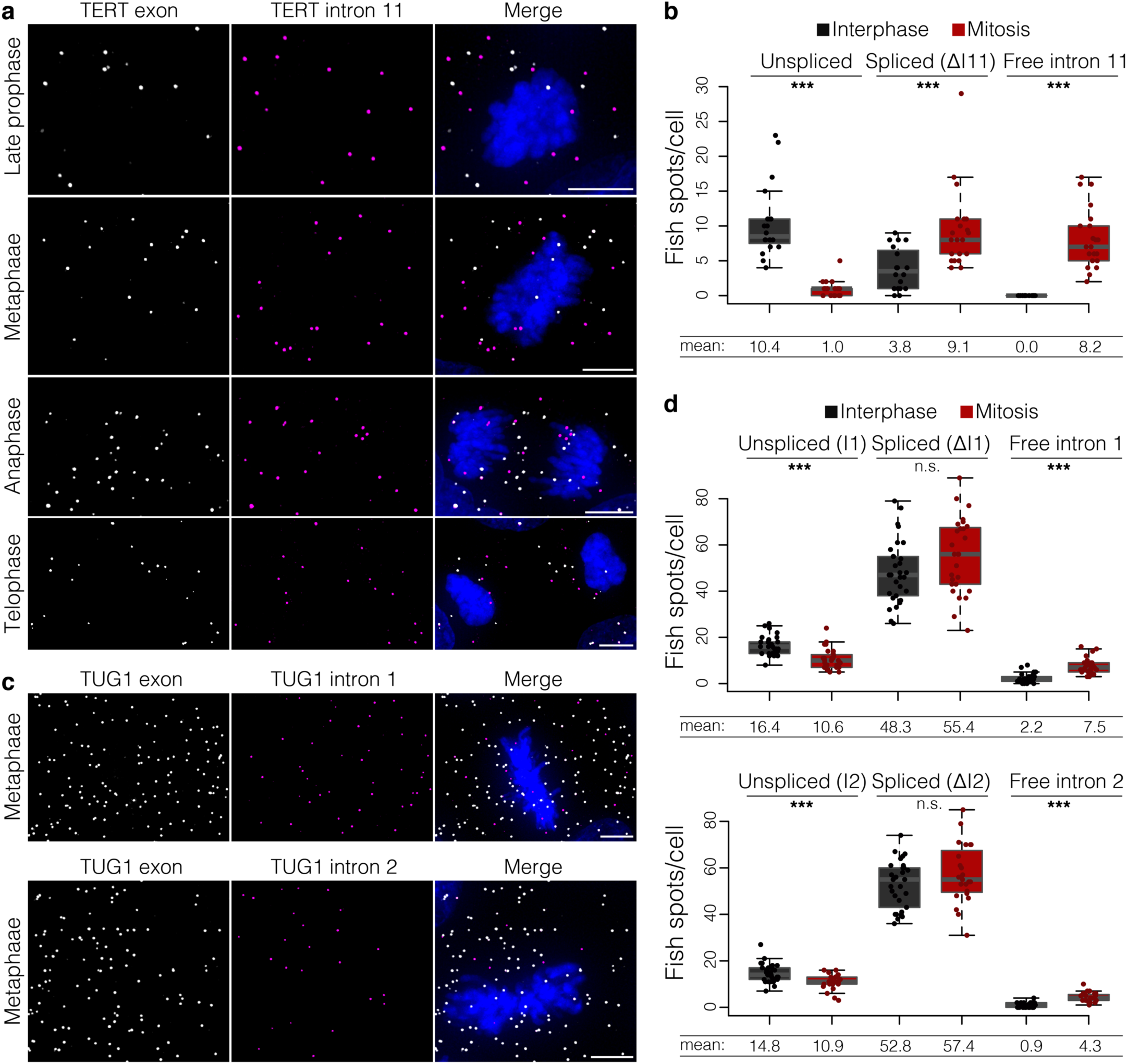
Splicing of TERT intron 11 occurs upon mitosis. **a**, Maximum intensity projections of TERT exon (gray) and intron 11 (magenta) smRNA FISH. Representative images of late prophase, metaphase, anaphase and telophase are shown. DAPI shown in blue. Scale bar, 5 μm. **b**, Quantification of unspliced TERT, spliced (ΔI11) TERT, and free intron 11 in interphase cells and during mitosis. \*\*\**P* ≤ 0.001, as evaluated by unpaired *t*-test versus interphase; *n* = 30 cells. **c**, Maximum intensity projections of TUG1 exon (gray) and intron 1 (magenta) or intron 2 (magenta) smRNA FISH. Representative images of metaphases are shown. DAPI shown in blue. Scale bar, 5 μm. **d**, Quantification of unspliced TUG1, spliced (ΔI1 or ΔI2) TUG1, and free intron 1 or 2 in interphase cells and during mitosis. n.s. = not significant, \*\*\**P* ≤ 0.001, as evaluated by unpaired *t*-test versus interphase; *n* = 30 cells.

TERT followed a similar trend, with nuclear, unspliced TERT RNA (assessed by retention of intron 11) being highly stable and retained in the nucleus even after 4.5 h of ActD treatment (Fig. 5c and Extended data Fig. 6b). Nuclear TERT was highly stable during the course of ActD treatment, while cytoplasmic TERT gradually decreased after transcription inhibition (∼4-fold decrease, *P*≤0.001, unpaired *t*-test for LN-18; ∼3-fold decrease, *P*≤0.001, unpaired *t*-test for iPS). Retention of intron 11 remained high (no significant decrease) for LN-18 and iPS, and the unspliced transcript remained in the nucleus.

As a control for transcription inhibition, we monitored GAPDH transcription sites visualized by smRNA FISH GAPDH exon/intron 2 overlap. GAPDH transcription sites were abolished in the majority of cells after 40 min of treatment, while after 2.5 and 4.5 h the signal was not detectable (Extended data Fig. 7). Collectively, these results show that intron-containing TUG1 and TERT are stable, long-lived transcripts, stably retained in the nucleus relative to their spliced cytoplasmic counterparts.

**Figure 7:**
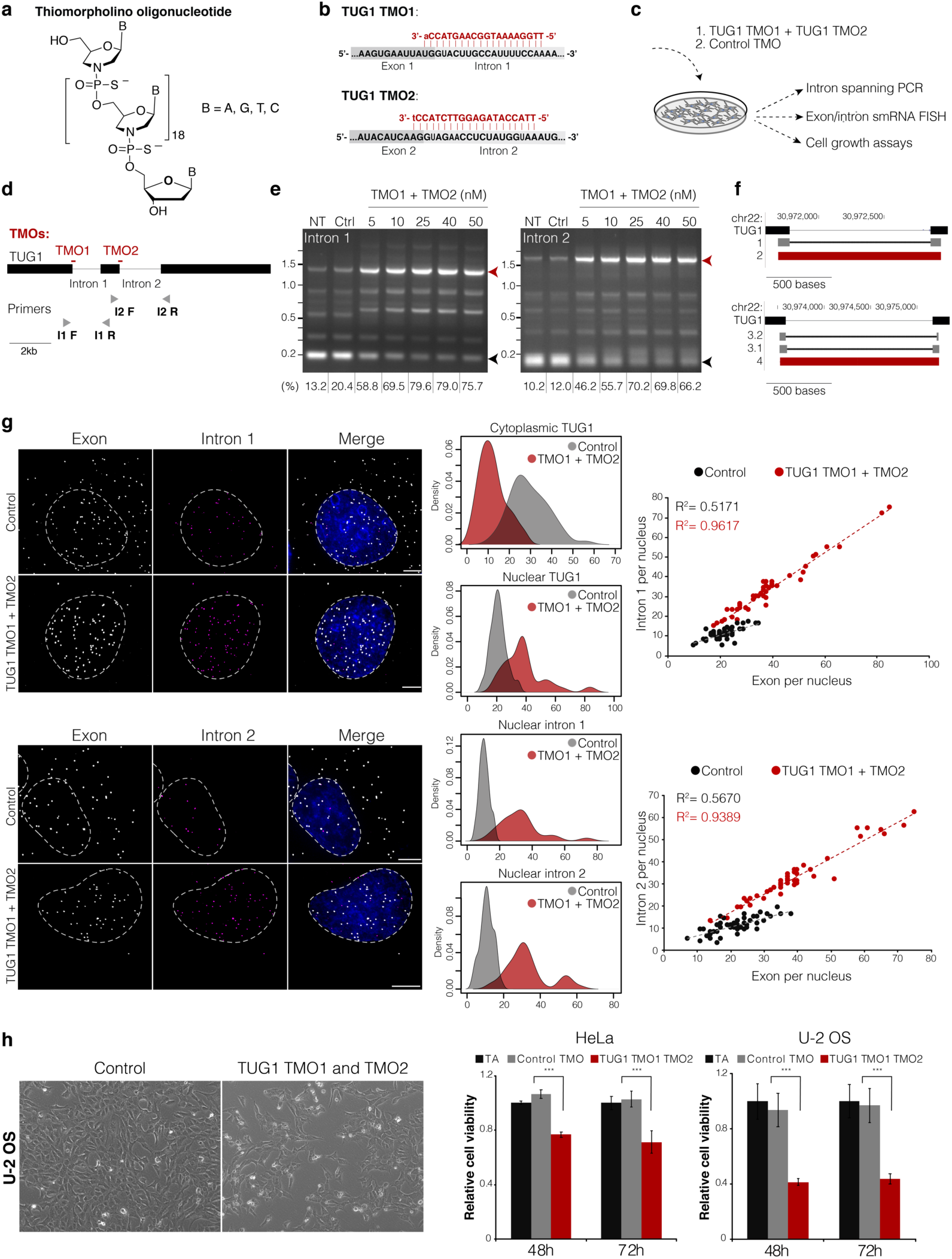
Intron retention drives nuclear compartmentalization of TUG1. **a**, The chemical structure of thiomorpholino oligonucleotide (TMO). **b**, The design of TUG1 TMO1 and TMO2 (in red) against the donor splice sites. For TMOs, upper-case red letters refer to thiomorpholino nucleotides and lower-case letters to 2’-deoxynucleosides at the 3’ end of each TMO. **c**, Experimental setup to assess the efficiency of TMO-based intron inclusion and its effect of subcellular localization of TUG1 and cell growth. **d**, TMO location scheme in respect to TUG1 transcript and the location on intron spanning primers (not to scale). **e**, PCR product of the intron spanning RT PCR of untreated (NT), control TMO (Ctrl) and increasing doses of a mixture of TUG1 TMO1 and TMO2. Black arrow, spliced product; red arrow, unspliced product. Below, the percentage of unspliced product. **f**, UCSC browser displaying Sanger sequencing results of spliced (band 1) and unspliced (band 2) products for intron 1 RT PCR (on top). Below, the sequences for spliced (band 3.1 and band 3.2) and unspliced (band 4) products for intron 2 RT PCR. **g**, Maximum intensity projections of TUG1 exon (gray) and intron 1 (magenta) or intron 2 (magenta) smRNA FISH in U-2 OS cells transfected with control TMO and with TUG1 TMO1 and TMO2. Nucleus in blue. Scale bar, 5 μm. Towards the right, distribution of nuclear TUG1, cytoplasmic TUG1, intron 1 or 2 retention in TUG1 TMO1 and TMO2 (red) versus control TMO (gray). **h**, Relative cell growth of HeLa and U-2 OS cells transfected with TUG1 TMO1 and TMO2, control TMO or transfection agent only (TA). Representative images of U-2 OS transfected with control TMO or TUG1 TMO1 and TMO2 shown on the left. \*\*\**P* ≤ 0.001, as evaluated by unpaired *t*-test versus control TMO; error bars represent SD; minimum three independent measurements.

### 5. TERT pre-mRNA splicing is cell-cycle specific occurring at mitosis

Interestingly, the above smRNA FISH analyses revealed that TERT pre-mRNA was spliced during cell division (after late prophase), when all TERT RNA molecules could be readily visualized as spliced; while TERT intron 11 was in the form of a solo intron, i.e., not co-localized with exons (Fig. 6a,b). The quantity of spliced TERT was increased in mitosis compared to interphase cells (mean value 9.1 vs 3.8 of spliced TERT molecules per mitotic or interphase cell, respectively, *P*≤0.001, unpaired *t*-test), while the quantity of unspliced TERT was reduced from a mean value of 10.4 molecules/cell in interphase cells to 1.0 in mitosis (*P*≤0.001, unpaired *t*-test). Lastly, while intron 11 was included in the vast majority of nuclear TERT mRNA in interphase cells, in mitosis the quantity of free intron 11 greatly increased (mean value 0.0 vs 8.2, respectively, *P*≤0.001, unpaired *t*-test). Importantly, the quantity of free intron 11 was comparable with the number of newly spliced TERT RNA (mean value 8.2 vs 9.1, respectively). Since the mitotically spliced intron was observed by smRNA FISH, it further indicates that the intron was stable, presumably in the form of a lariat. However, given that the intron lariat was not observed in other stages of the cell cycle, neither in the cytoplasm nor in the nucleus, it indicates that the stability of solo intron 11 is limited to mitosis.

We performed the same analysis for TUG1. In contrast to TERT, a great portion of TUG1 transcripts remained unspliced during mitosis compared to interphase cells (Fig. 6c,d, mean value 10.6 vs 16.4 for Δintron1, respectively; 10.9 vs 14.8 for Δintron 2, respectively), implying that TUG1 splicing is not dependent on mitosis. Together, our smRNA FISH analysis found that TERT splicing of retained intron 11 appears to be regulated in mitosis, opening an intriguing possibility of mitotic inheritance of fully spliced, cytoplasmic TERT mRNA.

### 6. Modified antisense oligonucleotides block splicing and affect subcellular RNA localization

Our observations of nuclear TERT and TUG1 intron retention are correlative and do not show a causality of intron retention driving their subcellular localization. To test the causality, we applied novel chemically modified antisense oligonucleotides (ASOs) called Thiomorpholinos (TMOs). TMOs are oligonucleotides having the bases (thymine, cytosine, adenine, and guanine) attached to morpholine, and these nucleosides are joined through thiophosphoramidate internucleotide linkages (Fig. 7a). They show increased hybridization stability towards complementary RNA (10°C increased melting temperature compared to an unmodified control duplex of identical sequence)^56^. TMOs are also highly stable towards exonuclease enzymes; minimal degradation is observed when treated with snake venom phosphodiesterase I for over 23 h. Unlike DNA:RNA duplexes, they do not elicit RNase H1 activity, making them ideal candidates for splicing studies.

We hypothesized that blocking excision of the retained introns of TUG1 and TERT would affect the subcellular localization of these RNAs, thereby opening potential novel avenues for RNA-directed therapies. TUG1 lncRNA was found upregulated in osteosarcoma primary samples and cell lines, where it was attributed to increased tumor proliferation and invasion, and its downregulation was shown to inhibit osteosarcoma proliferation *in vitro* and *in vivo*^48,57^. Furthermore, TUG1 was shown to be upregulated in cervical cancer and correlated with advanced clinical features and poor survival, where also TUG1 knockdown suppressed cervical cancer cell growth and metastasis *in vitro* and tumor growth *in vivo*^49^. Considering these findings, we reasoned that U-2 OS and HeLa cell lines would be good model cell lines to design the approach and study the effects of inhibiting TUG1 splicing.

First, we designed 20-mer TMOs against the two TUG1 donor splice sites, each hybridizing to 2 nt of the exon and 18 nt of the intron sequence (designated TUG1 TMO1 and TMO2) (Fig. 7b). To control for cell effects that could be caused by TMO intake, we designed a control TMO (randomized sequence of TMO1). TMOs were transfected at increasing concentration to U-2 OS and HeLa cells, after which cells were harvested for monitoring intron retention via intron-spanning RT-qPCR and smRNA FISH (Fig. 7c,d). The mixture of TUG1 TMO1 + TMO2 inhibited splicing and achieved retention of both introns in a dose-dependent manner already 24 h after treatment (Fig. 7e). Sanger sequencing of the spliced and unspliced RT-PCR products confirmed that the complete introns were retained (Fig. 6f).

We next used smRNA FISH to determine whether forced intron inclusion would affect the subcellular localization and availability of spliced TUG1 in the cytoplasm. Specifically, we performed dual color smRNA FISH in U-2 OS and HeLa cell lines treated with TUG1-targeting TMOs (TUG1 TMO1 + TMO2) and a control TMO. The TUG1-targeting TMOs gave a drastic shift in the subcellular localization and splicing of TUG1 (Fig. 7g and Extended data Fig. 8a). On average, in U-2 OS TUG1 decreased ∼2.4-fold in the cytoplasm (mean 29 in control vs 12 in TUG1 TMO1+2), while TUG1 increased ∼1.8-fold in the nucleus (mean 21 in control vs 38 in TUG1 TMO1+2). Similarly, in HeLa cells TUG1 decreased ∼2.7-fold in the cytoplasm (mean 22 in control vs 8 in TUG1 TMO1+2), and it increased ∼1.7-fold in the nucleus (mean count 29 in control vs 48 in TUG1 TMO1+2) (Fig. 7g and Extended data Fig. 8a). After TUG1 TMO1 + TMO2 application, intron retention in nuclear TUG1 was significantly increased in U-2 OS and HeLa cells (PIR intron 1 increased from 51% to 85%, PIR intron 2 increased from 52% to 84% in U2-OS; PIR intron 1 increased from 67% to 92%, PIR intron 2 increased from 57% to 92% in HeLa).

**Figure 8:**
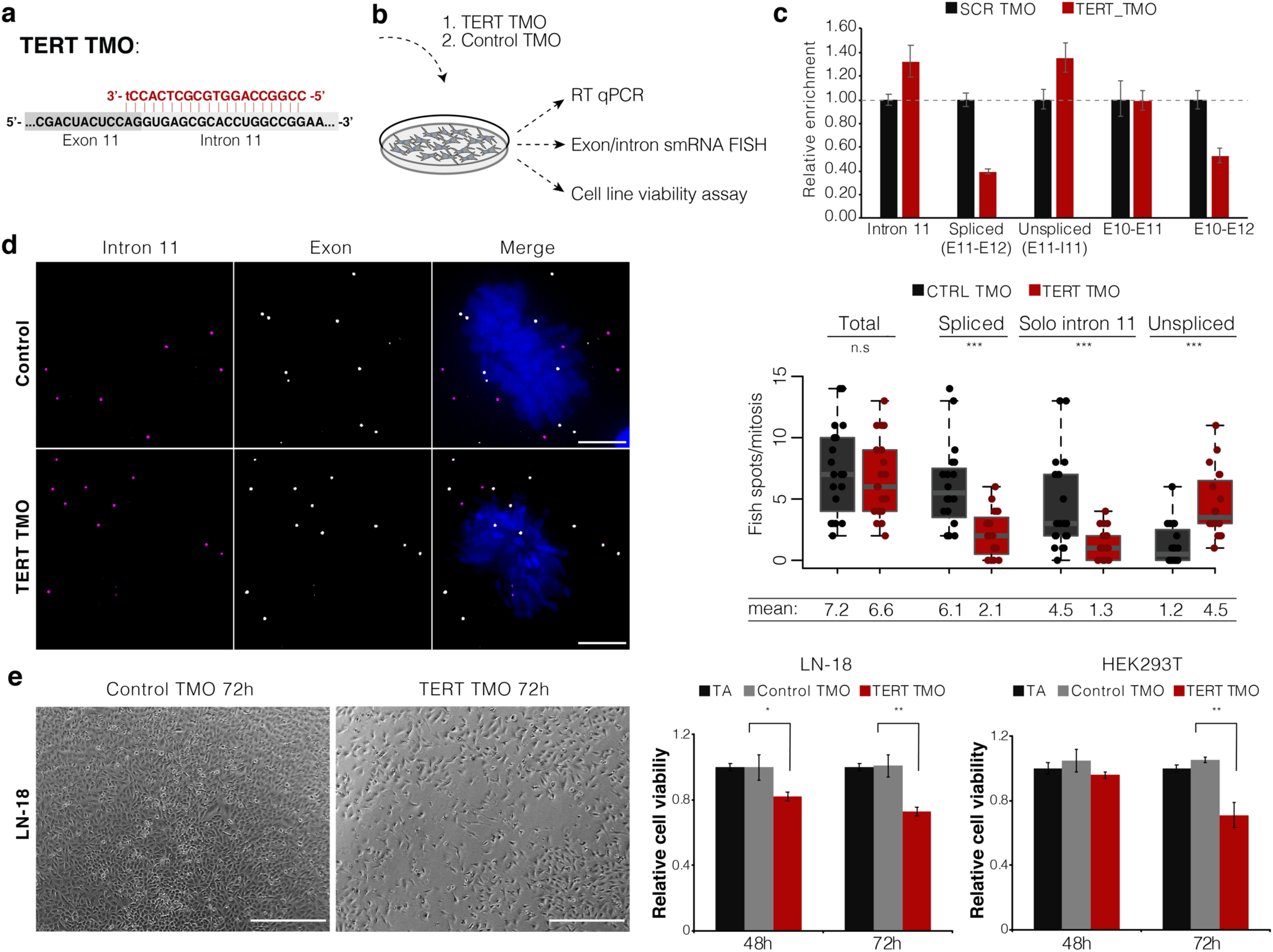
TMO-based prevention of TERT splicing reduces cell growth *in vitro*. **a**, Scheme showing the design of TERT TMO (in red) against the exon11/intron11 donor splice site. The upper-case red letters refer to thiomorpholino nucleotides and the lower-case letter to a 2’-deoxynucleoside at the 3’ end. **b**, Experimental setup to assess the efficiency of TMO-based TERT intron 11 inclusion (RT qPCR and smRNA FISH) and its effect on cell growth. **c**, Relative expression of TERT intron 11, spliced TERT (Exon10-Exon11, Exon10-Exon12, Exon11-Exon12), and unspliced TERT (Exon11-Intron11) over GAPDH assessed by RT qPCR. Error bars represent SD, three replicates. **d**, Maximum intensity projections of TERT exon (gray) and intron 11 (magenta) smRNA FISH in LN-18 cells transfected with control TMO and TERT TMO. DAPI, blue. Scale bar, 5 μm. On the right, quantification of total TERT (exon signal), unspliced TERT, spliced TERT (ΔI11), and free intron 11 during mitosis of LN-18 cells transfected with control TMO (CTRL) or TERT TMO. n.s. = not significant, \*\*\**P* ≤ 0.001, as evaluated by unpaired *t*-test versus control TMO; *n* = 30 cells. **e**, Cell growth of LN-18 and HEK293T cells transfected with TERT TMO, control TMO or transfection agent only (TA). Representative images of LN-18 transfected with control TMO or TERT TMO shown on the left. Scale bar, 25 μm. \**P* ≤ 0.05, \*\**P* ≤ 0.01, as evaluated by unpaired *t*-test versus control TMO; error bars represent SD; minimum two (HEK293T), three (LN-18) independent measurements.

In parallel, we used TUG1 TMO1 labelled with FITC (TUG1 TMO1-FITC) to determine the subcellular localization of the TMO after transfection. TUG1 TMO1-FITC showed predominantly nuclear localization, and it was stably localized in the nucleus 96 h after transfection (later time points were not assessed) (Extended data Fig. 8b), consistent with these oligos being able to alter nuclear splicing processes. Together, our results demonstrate that TMOs can be used to achieve increased intron retention and in turn increased nuclear localization of transcripts. We note that this would have numerous applications for rendering active cytoplasmic transcripts as inactive in the nucleus (i.e., preventing translation).

### 7. Functional consequences of enforced intron retention for TUG1 and TERT

Having observed that increasing intron retention increases nuclear localization of TUG1, we wanted to determine if this redistribution of transcripts has a functional cellular consequence. Specifically, U-2 OS and HeLa cell lines were transfected with 25 nM of TUG1 TMO1 + TMO2 and cell growth was assessed 48 h and 72 h post-transfection relative to transfection agent only and a control TMO (Fig. 7h). Both cell lines showed a reduction in cell growth after 48 h of TUG1 TMO treatment compared to controls (mean 24% and 59% reduced growth for HeLa and U-2 OS, respectively, *P*≤0.01, unpaired *t*-test), and after 72 h (mean 29% and 57% reduced growth for HeLa and U-2 OS, respectively, *P*≤0.01, unpaired *t*-test). Thus, in both cases altering the subcellular distribution of TUG1 impaired cell growth.

To determine if our TMO strategy is also applicable to pre-mRNAs we focused on TERT. Briefly, we designed a TMO to retain specified intron 11 of TERT and determine the cellular consequences thereof. To this end, we synthesized a 20-mer TMO targeting TERT exon 11/ intron 11 junction (Fig. 8a). Cell lines with uniform reactivation of TERT expression, LN-18 and HEK293T, were transfected with TERT TMO and control TMO. Because TERT intron 11 is quite long (3.8 kb), intron-spanning PCR was not feasible to assess intron retention. Thus, we applied exon/intron junction RT-qPCR to assess the efficiency of intron retention (Fig. 8b,c). We found that TMOs enforcing intron 11 retention decreased splicing of intron 11 by ∼60% compared to the control TMO. In contrast, intron 11-containing TERT (assessed by monitoring exon 11 to intron 11 junction) was increased ∼35% compared to control TMO. As additional controls, we applied primers at the upstream exon 10 to exon 11 junction, which was not affected with TERT TMO treatment, and exon 10 to exon 12 junction, which was decreased ∼50%, in accordance with the decrease in exon 11 to exon 12 junction (Fig. 8c).

We further leveraged the specific retention of intron 11 and the restriction of splicing to mitosis. We observed that the total number of TERT RNA molecules (assessed by overall exon signal) was not altered during mitosis between control and TERT TMO (mean values 7.2 and 6.6, respectively). Consistent with the above results, we observed a significant effect of TERT TMO on splicing of intron 11 only during mitosis (Fig. 8d). More specifically, the majority of intron 11 was spliced out and observed in the form of a solo intron with control TMO, and these solo introns were significantly decreased in cells treated with TERT TMO (mean value 4.5 solo intron 11/mitosis in control TMO and 1.3 solo intron 11/mitosis in TERT TMO, *P*≤0.001, unpaired *t*-test). The observation of solo intron 11 during mitosis is in accordance with the observation made in iPS cells (Fig. 5a,b). While during mitosis most of TERT RNA was in the form of spliced RNA in the control TMO, the quantity of spliced TERT significantly decreased in TERT TMO samples (mean values 6.1 and 2.1, respectively, *P*≤ 0.001, unpaired *t*-test). Consequently, the quantity of unspliced TERT increased in TERT TMO compared to control TMO samples (mean values 4.5 and 1.2, respectively, *P*≤0.001, unpaired *t*-test). Overall, the RT-qPCR and smRNA FISH confirmed that TMOs can specifically inhibit splicing of intron 11 from TERT pre-mRNA. We next sought out to determine whether inhibiting the availability of spliced TERT by TERT TMO would affect cell growth of LN-18 and HEK293T cell lines. Both cell lines were transfected with 25 nM of TERT TMO, and cell proliferation was assessed (Fig. 8e). LN-18 cell line showed a reduction in cell growth after 48 h of treatment compared to transfection agent only and control TMO (mean reduction of 18%, *P*≤0.05), which was further enhanced after 72 h (mean reduction of 28% in cell growth, *P*≤0.01). HEK293T showed a delayed response and cell growth was reduced after 72 h treatment with TERT TMO (mean reduction of 29% in cell growth, *P*≤0.01) compared to control TMO and transfection agent only. It is intriguing that cell growth is compromised so quickly after reduction of translatable TERT mRNA, given that telomere shrinkage due to inhibition of telomerase typically takes many population doublings before it gives a growth defect^58^.

Collectively, these findings demonstrate that TMOs effectively block splicing and change cellular localization and availability of the RNA. Moreover, we find that this perturbed subcellular transcript distribution has a functional consequence on cell growth.

## Discussion

It has long been known that the spatio-temporal distribution and compartmentalization of RNA in the cell is tightly coupled with its subcellular function^1–6^. Studies of underlying mechanisms have pinpointed RNA motifs and structural features that target RNA subcellular localization^59,60^. Splicing has been shown to strongly influence RNA localization^9,10^. For example, lncRNAs are inefficiently spliced, display increased intron retention relative to coding mRNAs, and are more nuclear than mRNAs^11,61,62^. Despite these intriguing findings, the causality of splicing events, such as intron retention, in the molecular events driving nuclear retention of RNAs has remained unclear. A surprising finding is that the majority of TERT transcripts are nuclear, and therefore translationally inert, yet the underlying mechanism remained unknown^39,40^.

Here we addressed this question primarily using single molecule RNA FISH to spatio-temporally measure specific splicing events that may alter subcellular RNA localization. We focused on two cancer-related transcripts, TERT mRNA and TUG1 lncRNA. We find that both mRNA and lncRNA localization patterns are driven by consistent retention of specific introns. It has been shown that some nuclear-retained, stable, intron-retained RNAs are poised or ‘detained’ for a signal for post-transcriptional splicing, hence serving as a reservoir of RNAs readily available depending on cellular activity^12,13^. In this regard, the striking splicing of retained TERT intron 11 after cells’ entry to mitosis was an intriguing indication that fully spliced TERT might be generated mitotically. Retention of specific introns would compartmentalize TERT RNA in the nucleus of interphase cells, while upon cells’ entry to mitosis, retained introns would be spliced out and daughter cells would inherit fully spliced TERT.

Mitotic inheritance of spliced TERT would ensure that telomere elongation occurs only in mitotically active cells, still allowing telomerase assembly during the later stages of the cell cycle when DNA is replicated and telomeres elongated^63–65^. Together, these findings raise the question of how intron 11 is retained in order to specifically be spliced out during mitosis and produce a cytoplasmic transcript for translation. In this regard it is interesting to consider that TERT intron retention may be regulated as part of a broader program of differential intron retention (and other forms of alternative splicing) that is controlled by the SR protein splicing factor kinase CLK1 during the cell cycle^66^. Alternatively, possibly some other signaling pathway could regulate the splicing of the retained introns and nuclear export of TERT for translation during interphase.

We found that the lncRNA TUG1 is equally distributed between nucleus and cytoplasm across multiple cell lines. Hence the same locus gives rise to equal amounts of either efficiently spliced cytoplasmic TUG1 or intron-retained nuclear TUG1, where intron retention dictates nuclear/cytoplasmic transcript distribution. This interesting splicing balance could have important implications: (i) the longer TUG1 lncRNA with retained introns could exert a specific nuclear RNA function in this longer form, which is consistent with the strong conservation of TUG1 intronic sequences; (ii) intron sequences could give rise to distinct functions; (iii) the efficiently spliced cytoplasmic TUG1 could be destined to encode a protein, as has been proposed by recent studies^51^; (iv) the conserved distribution of TUG1 in the nucleus and cytoplasm could represent a translational buffering or two distinct functionalities. One of these mechanisms, or their combination, potentially underlies a 100% penetrant male infertility phenotype in TUG1 knock-out mouse models^51^.

Both TERT and TUG1 are upregulated in many cancers and thus represent important therapeutic targets. To this end, we tested a novel RNA-based strategy to alter TERT and TUG1 splicing and subcellular distribution. We found that this TMO antisense approach was highly effective and specific at blocking TERT and TUG1 splicing events. Importantly, altering these specific splicing patterns using our TMO approach not only affected subcellular distribution but, in both cases, affected cell growth. Thus, TMO-based strategies could be universally applicable not only to other transcripts that retain specific introns, but to a variety of oncogene transcripts that could be rendered inert in the nucleus.

## Materials and Methods

### Cell lines and cell culture

Cell lines were obtained from ATCC and cultured according to recommended protocols. Human iPS WT-11 cells were cultured on Vitronectin (Thermo Fisher Scientific) coated 6-well plates or glass coverslips (for smRNA FISH purposes) in Essential 8 Flex medium (Thermo Fisher Scientific) with E8 supplement (Thermo Fisher Scientific), Rock inhibitor and 2.5% penicillin-streptomycin. iPS cells we passaged with EDTA in dPBS. Mouse embryonic stem cells were cultured on top of gelatin (0.1%, EMD Millipore) coated plates or glass coverslips (for smRNA FISH purposes). Embryonic stem cell media was prepared as follows: KnockOut DMEM medium (Thermo Fisher) supplemented with ESC FCS (Millipore Sigma), non-essential amino acids (Thermo Fisher), GlutaMAX supplement (Thermo Fisher), penicillin-streptomycin (Thermo Fisher), 50 mM 2-mercaptoethanol LIF and 2i (and CHIR99021, Sigma-Aldrich and PD0325901, Sigma-Aldrich).

Actinomycin D (Sigma-Aldrich) was used at final concentration of 5 μg/mL in full growth media. Cell pellets and coverslips were harvested at 0, 40 min, 2.5 h and 4.5 h after adding Actinomycin D, and processed for RNA extraction and smRNA FISH as described below.

### RNA extraction

After the corresponding treatments, cell pellets were harvested and RNA extraction was performed with Maxwell LEV Simply RNA tissue kit (Promega) following manufacturer’s instructions with DNase I treatment. Each sample was tested for DNA contamination by qPCR after each extraction. RNA quality was assessed on 2% agarose gel and Bioanalyzer (RNA Nano Assay: 25–500 ng/μL).

### Analysis of intron retention from RNA-seq

Vast-tools v2.2.2^53^ (https://github.com/vastgroup/vast-tools) was used to calculate PIR values from human iPS cells as well as mouse ES cells and published data from iPS cells^67^ (GEO: GSE42100). Reads mapping to mid-intron sequences and balanced counts of reads aligning to upstream and downstream exon-intron sequences were used to evaluate intron retention levels. PIR was measured as a percentage of mean retention reads over the sum of retained and spliced intron reads. Raw values were filtered based on reported quality scores, requiring at least 15 total reads per event and absence of a positive result (*P*<0.05) for the binomial test for upstream/downstream junction read balance. PIR values for human TERT intron 11 were reported by vast-tools as imbalanced due to an alternative exon within the intron and were therefore re-calculated based solely on the downstream intron-exon junction reads. Similarly, TUG1 intron 1 was re-calculated based on upstream exon-intron junction reads due to an alternative acceptor site, and TUG1 intron 2 was absent from the VastDB database. For the analysis of global levels of maximum and minimum PIR in coding and non-coding genes, gene biotype annotations were taken from GENCODE v29 (human) and vM23 (mouse) and simplified to ‘coding’, ‘lncRNA’, and ‘other’ (not shown).

### cDNA synthesis and qPCR analysis

Reverse transcription was performed with SuperScript® IV First-Strand Synthesis System (Thermo Fisher Scientific) with Superase RNase inhibitor (Ambion) and random hexamers on 0.2–1 μg of RNA. Relative expression was determined by qPCR using SYBR Green I master mix (Thermo Fisher Scientific) according to manufacturer’s instructions using the following amplification conditions: 95°C 10′; 45 cycles of 95°C 15″, 57.5°C 20″ and 72°C 25″. Expression levels were normalized using GAPDH. A list of primers used in qPCR analyses are summarized below. Their efficiencies were compared to ensure analysis by the comparative Ct method. Relative expression data was analyzed comparing the Ct values of the gene of interest with Ct values of the reference gene for every sample. We used the formula 2ΔΔCt, ΔΔCt being the difference between the Ct of the RNA of interest and the Ct of the housekeeping gene. Duplicates or triplicates were made for each sample and primer set.

### RT PCR and Sanger sequencing

cDNA was amplified with primers listed in Primers and sequences section of methods with Q5® High-Fidelity DNA Polymerase PCR System (NEB). PCR conditions: initial denaturing at 95°C 2’; 40 cycles of denaturing at 95°C 30’’, annealing at 58°C 30’’ and extension at 72°C 2’ 30’’; followed by final extension at 72°C 7’. PCR product was examined on a 1% agarose gel for correct size and specificity. Bands corresponding spliced or unspliced TUG1 were cut from the gel and DNA was extracted with Gel extraction kit (Qiagen) according to instructions. Extracted DNA was cloned with TOPO PCR Cloning Kit (Thermo Fisher Scientific), and positive colonies selected on ampicillin agar plates. Minipreps from ∼5 colonies for each amplicon were sent for Sanger sequencing to Genewiz using T3 or T7 primers.

### Single molecule RNA FISH

smFISH was performed as previously described^30^. Tiled oligonucleotides targeting human and mouse TUG1 exons, TERT intron 2, TERT exons, GAPDH intron 2 and GAPDH exons labeled with either Quasar 570 or Quasar 670 were used in our previous studies^40,51^. For this study, we custom designed tiled oligonucleotides targeting human and mouse TUG1 intron 1 (Quasar 570) and intron 2 (Quasar 570), TERT intron 11 (Quasar 670) and TERT intron 14 (Quasar 670) using LGC Biosearch Technologies’ Stellaris online RNA FISH probe designer (Stellaris Probe Designer, version 4.2) which were produced by LGC Biosearch Technologies. Cells were seeded on glass coverslips coated with poly-L-lysine (10 μg/mL in PBS), vitronectin (human iPS cells) or gelatine (mouse ES cells). Coverslips were washed 2 times with PBS, fixed in 3.7% formaldehyde in PBS for 10 min at room temperature (RT), followed by washing 2 times with PBS and immersed in 70% EtOH at 4°C for a minimum of 1 h. Prior hybridization, coverslips were washed with 2 mL of wash buffer A (LGC Biosearch Technologies) supplemented with 10% deionized formamide (Agilent) at RT for 5 min. Cells were hybridized with 80 μL of hybridization buffer (LGC Biosearch Technologies) supplemented with 10% deionized formamide (Agilent) containing 1:100 dilution of smRNA FISH probes overnight at 37°C in a humid chamber. The next day, cells were washed with 1 mL of wash buffer A with 10% formamide for 30 min at 37°C, followed by a wash with wash buffer A with 10% formamide containing Hoechst DNA stain (1:1,000; Thermo Fisher Scientific) for 30 min at 37°C. Coverslips were washed with 1 mL of wash buffer B (LGC Biosearch Technologies) for 5 min at RT, equilibrated 5 min in base glucose buffer (2x SSC, 0.4% glucose solution, 20 mM Tris pH 8.0 in RNase-free H_2_O), and then incubated 5 min in Base Glucose buffer supplemented with 1:100 dilution of glucose oxidase (stock 3.7 mg/mL) and catalase (stock 4 mg/mL). Afterwards, the coverslips were mounted with ProlongGold or ProlongGlass (Life Technologies) on a glass slide and left to curate overnight before proceeding to image acquisition (see below).

### Microscopy and image analysis

Z stacks with 200-250 nm z-step capturing the entire cell volume were acquired with a GE wide-field DeltaVision Elite microscope with an Olympus UPlanSApo 100×/1.40-numerical aperture oil objective lens and a PCO Edge sCMOS camera using appropriate filters. The built-in DeltaVision SoftWoRx Imaging software was used to deconvolve the three-dimensional stacks. Maximum intensity projections were generated in Fiji and subjected for quantification using Fiji. The brightness and contrast of each channel was adjusted. Overlapping exon/intron spots were considered as intron-retained transcripts, while exon only transcripts as spliced transcripts. Each imaging experiment was performed at least two times quantifying at least 50 cells across independently acquired datasets. For ActD treatment and mitosis, less cells/mitosis were quantified per treatment, as indicated in the figure legend. Analysis of z-stacked was additionally performed in 3D in Imaris to confirm that nuclear intron-retained transcripts were within the nucleus.

### TMO synthesis

Prior to thiomorpholino oligonucleotide (TMO) synthesis, appropriately protected morpholino nucleosides of adenine, guanine, thymine and cytosine and their corresponding phosphorodiamidites were synthesized as reported elsewhere^56^. All TMOs were synthesized using an Applied Biosystems Model 394 Automated DNA Synthesizer using conventional DNA synthesis reagents that were purchased from Glen Research, VA. Briefly, 1.0 μM succinyl CPG support was detritylated using 3% trichloroacetic acid in dichloromethane. The 5’-unprotected nucleoside was allowed to react with a 1.0 M solution of the appropriate morpholinonucleoside phosphorodiamidite in acetonitrile in the presence of 0.12 M 5-ethylthio-1H-tetrazole (600s coupling time). After sulfurization using 0.05 M sulfurizing reagent II in pyridine/acetonitrile, the capping step was carried using conventional Cap Mix A (acetic anhydride/tetrahydrofuran) and Cap Mix B (1-methylimidazole in acetonitrile), completing one synthesis cycle. Multiple synthesis cycles were repeated until a TMO oligonucleotide of the desired sequence was obtained. The 5’-DMT group on the solid-support bound final oligonucleotide was not detritylated so that purification could be carried out using the DMT-On/Off procedure^68^.

Cleavage and deprotection was carried out using 28% aqueous ammonia at 55°C for 16 h. After cooling to 25°C followed by evaporation of the ammonia mixture, the oligonucleotides were purified by ion-pair reversed phase HPLC. During this process, the total reaction mixture (after evaporation to dryness) was dissolved in 3% aqueous acetonitrile and injected into an Agilent 1100 HPLC equipped with a manual injector. Due to the lipophilicity of the DMT handle, the DMT-On TMO oligonucleotide could be easily separated from failure products using a gradient of 50 mM Triethylammonium bicarbonate in acetonitrile (Agilent Zorbax C18 column, 2.0 mL flow rate). The DMT-On fractions were pooled, evaporated to dryness and treated with 50% aqueous acetic acid for 5 min. After quenching with triethylamine, the mixture was evaporated to dryness. The resulting solids were dissolved in 3% aqueous acetonitrile and the deprotected TMO oligonucleotides were re-purified by ion-pair RP-HPLC. All oligonucleotides were desalted prior to use. Graphical illustration of thiomorpholino oligonucleotide synthesis shown in Extended data Fig. 9.

### Transfection and TMO treatment

Cells were plated at 200,000 cells/well in a 6-well plate, or 100,000 cells/well in a 12-well plate, the day prior to transfection. Each cell line was transfected with increasing quantity of TMOs with two different transfection agents (Lipofectamine RNAiMAX (Thermo Fisher Scientific) and Xtreme Gene siRNA transfection agent (Sigma)) to determine the optimal transfection conditions for each cell line. Fluorescently labeled TMO was used to assess transfection efficiency, while intron-spanning PCR (only for TUG1), RT-qPCR and smRNA FISH were used to assess the efficiency of intron inclusion. Lipid-oligo complexes were prepared at room temperature in OptiMem medium (Thermo Fisher Scientific) according to the manufacturer’s instructions. After incubation time, lipid-oligo complexes were added dropwise to wells containing freshly added full growth media. U-2 OS was most efficiently transfected with Xtreme Gene, while HeLa, LN-18 and HEK cell lines were more efficiently transfected with Lipofectamine RNAiMAX. 25 nM TMO was chosen as the lowest quantity achieving maximum intron inclusion efficiency.

### Cell growth assays

Cells were plated at density of 1,000 cells/well in a 96 well plate. After 24 h, cells were transfected with 25 nM of the corresponding TMO. 48 h and 72 h post-transfection, cell culture media was replaced by 10% of AlamarBlue reagent (DAL1100, ThermoFisher Scientific) in full growth media 2-4 h prior to reading fluorescence. Fluorescent data was collected using the CLARIOstar microplate reader from BMG Labtech fluorescence plate reader following the manufacturer’s recommendations.

### TUG1 and TERT conservation analysis

Human and mouse TUG1 and TERT genomic sequences were downloaded from hg38 and mm10, respectively. Alignments were prepared in Geneious using MAFFT v7.388^69,70^. Alignments were imported in CLC main workbench (Qiagen) where sequence conservation was further analyzed by pairwise sequence comparison and visualized.

### Splice site strength analysis

MaxEntScan^71^ was used to calculate maximum entropy scores for 9 nt donor splice sites and 20 nt acceptor splice sites.

### RNA sequencing and read alignment

RNA from U-2 OS, HeLa and mES cell lines was extracted with Maxwell LEV Simply RNA isolation kit. RNA quality was assessed with BioAnalyzer. 1.5 μg of total RNA was sent to Novogene for library preparation and sequencing. Poly(A) RNA enrichment and library preparation was performed with NEBNext® Poly(A) mRNA Magnetic Isolation Module (NEB E7490) and NEBNext® Ultra RNA Library Prep Kit for Illumina® (E7530), and sequenced on the Novaseq6000. We retrieved RNA-seq data for HEK293, LN-18, HCT116 and fibroblasts (accession numbers: SRR3997506, SRR8769945, SRR8615282, SRR5420980) and gene annotations were retrieved from Gencode (vM23*). Raw reads were mapped to GRCm38 using the NF-CORE RNA-seq pipeline (v1.4.2*)^72^.

### Data availability

Browser tracks can be found at: https://genome.ucsc.edu/s/GabrijelaD/TERT_multiple_cell_lines_Share (HEK293, LN-18, HCT116, HeLa and BJ fibroblasts for TERT); https://genome.ucsc.edu/s/GabrijelaD/TUG1_multiple_cell_lines_Share (U2-OS, HeLa, BJ fibroblasts, LN-18, HEK292, HCT116 for TUG1).

## SEQUENCES AND PRIMERS

### TMO sequences

**TUG1 TMO1:** 5’-TTGGAAAATGGCAAGTACCA-3’

**TUG1 TMO2:** 5’-TTACCATAGAGGTTCTACCT-3’

**TERT TMO:** 5’-CCGGCCAGGTGCGCTCACCT-3’

**Control TMO:** 5’-ACACGGATATCGGTAAGAAT-3’

### TUG1 primers

**TUG1 exon 2 F:** 5’-AGCCTTCAGAGACACACAATAA-3’

**TUG1 exon 2 R:** 5’-TCCAAAGAAGATGCTATGAGGAG-3’

**TUG1 intron 1 F:** 5’-AAGGCATTGGAAGAGGAAGAG-3’

**TUG1 intron 1 R:** 5’-CTGGCTTAGGCAAAGACAAATG-3’

**TUG1 intron 2 F:** 5’-GGTATTGGGAACCTCAGGAAAT-3’

**TUG1 intron 2 R:** 5’-GGCCCAGGAATATCAGTAAGTC-3’

### TUG1 intron spanning primers

**TUG1 exon 1 F (for splicing):** 5’-CCAGCACTGTTACTGGGAATTA-3’

**TUG1 exon 2 R (for splicing):** 5’-GGTCTGTAGGCTGATGGAATAG-3’

**TUG1 exon 2 F (for splicing):** 5’-CCCTTACCTAACAGCATCTCAC-3’

**TUG1 exon 3 R (for splicing):** 5’-TCACTCAAAGGGCTTCATGG-3’

### TERT primers

**TERT exon 10 F:** 5’-CTCCTGCGTTTGGTGGATGA-3’

**TERT exon 11 R:** 5’-AAGTTCACCACGCAGCCATA-3’

**TERT exon 11 F:** 5’-GTCCGAGGTGTCCCTGAGTAT-3’

**TERT exon 12 R:** 5’-TGTGACACTTCAGCCGCAA-3’

**TERT intron 11 F:** 5’-GCCAATCCCAAAGGGTCAGA-3’

**TERT intron 11 R:** 5’-TCGGGTTCAGAGGGACTCAT-3’

**TERT intron 14 F:** 5’-GAGCAGAGCACCTGATGGAA-3’

**TERT intron 14 R:** 5’-GGCTCTGTCGTGGTGATACG-3’

### GAPDH primers

**GAPDH F:** 5’-GGAGCGAGATCCCTCCAAAAT-3’

**GAPDH R:** 5’-GGCTGTTGTCATACTTCTCATGG-3’

**GAPDH intron F:** 5’-AGGTCCTCTTGTGTCCCCTC-3’

**GAPDH intron R:** 5’-TTCCAACTACCCATGACTCAGC-3’

### 45s rRNA

**45S rRNA F:** 5’-TGTCAGGCGTTCTCGTCTC-3’

**45S rRNA R:** 5’-AGCACGACGTCACCACATC-3’

## Acknowledgments

We thank the Rinn lab members for insightful discussions. We thank Roy Parker and Carolyn Decker (University of Colorado Boulder) for access to the DeltaVision Elite microscope. We thank Theresa Nahreini and Nicole Kethley for use of the Cell Culture Facility (University of Colorado Boulder). iPS and LN-18 cells were a generous gift from Teisha Rowland from the Stem Cell Research and Technology Resource Center (University of Colorado Boulder). U.B. is supported by funds from a Canadian Institutes for Health Research Foundation Grant to B.B.. T.R.C. is an investigator of the Howard Hughes Medical Institute (HHMI). J.L.R. is an HHMI Faculty Scholar and holds a Marvin H. Caruthers Endowed Chair for Early Career Faculty. This research was supported by NIH grant P01GM099117 to J.L.R.

## Author contributions

Study conceptualization and design: G.D., T.R.C., M.C. and J.L.R.; experiment design, performance, microscopy and data analysis: G.D.; TMO synthesis: H.K., K.J..; computational analyses: G.D., U.B., M.S.; intellectual input: G.D., U.B., H.K., B.B., T.R.C., M.C. and J.L.R.; funding: M.C. and J.L.R.; writing the paper: G.D. and J.L.R. with input from all of the authors.

