## Extended data Figures for "Nuclear compartmentalization of TERT mRNA and TUG1 lncRNA transcripts is driven by intron retention: implications for RNA-directed therapies"

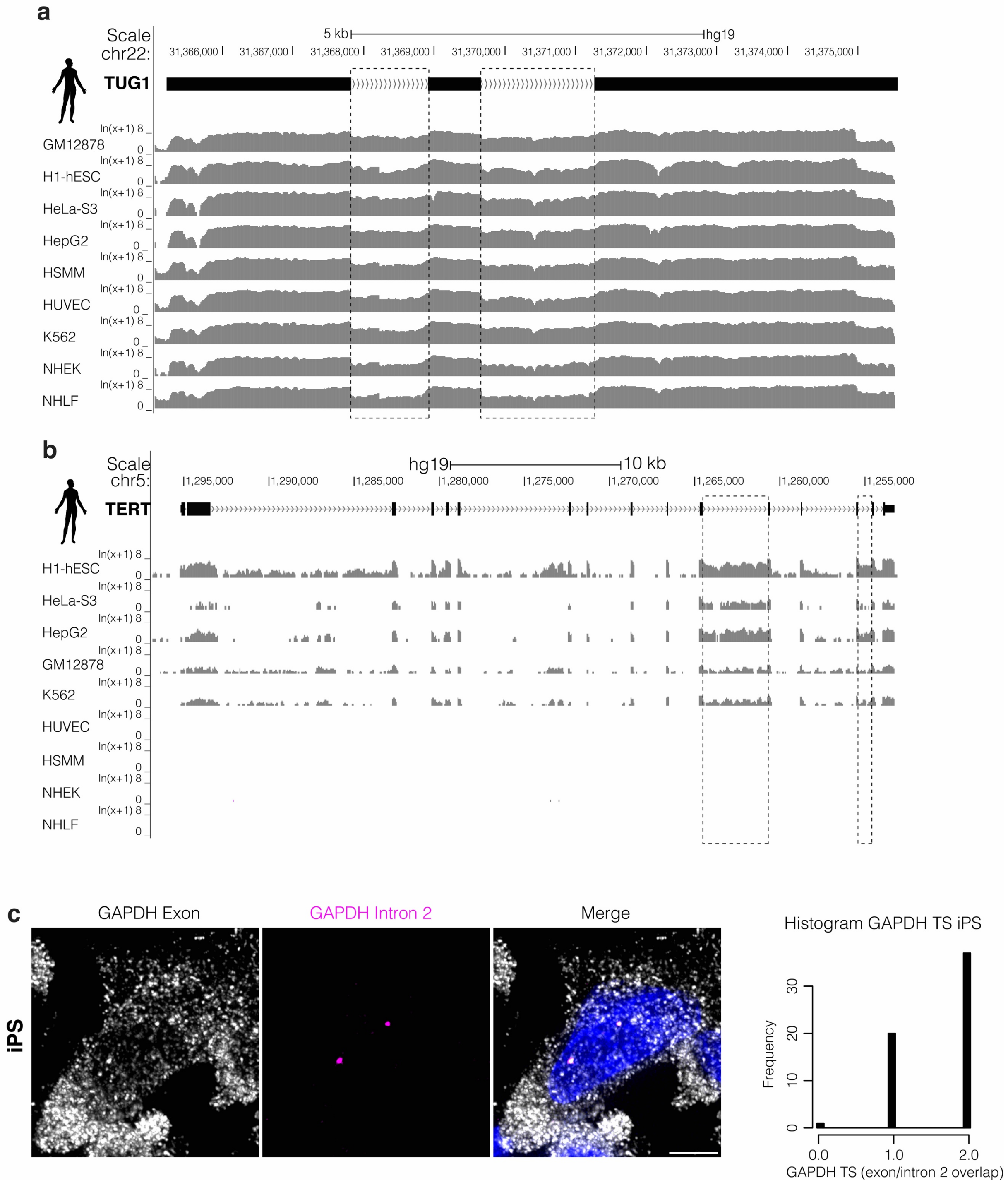


**Extended data figure 1: a**, UCSC Genome Browser showing RNA-seq tracks across the TUG1 locus from various cell lines available from ENCODE are depicted. **b**, UCSC Genome Browser showing RNA-seq tracks across the TERT locus from various cell lines available from ENCODE are depicted **c**, Maximum intensity projection of GAPDH exon (gray) and intron 2 (magenta) smRNA FISH on iPS cells. Nucleus in blue. Scale bar, 5 µm. On the right, histogram showing the number of exon/intron2 signal overlap in iPS cells.

*Extended Data Figures*

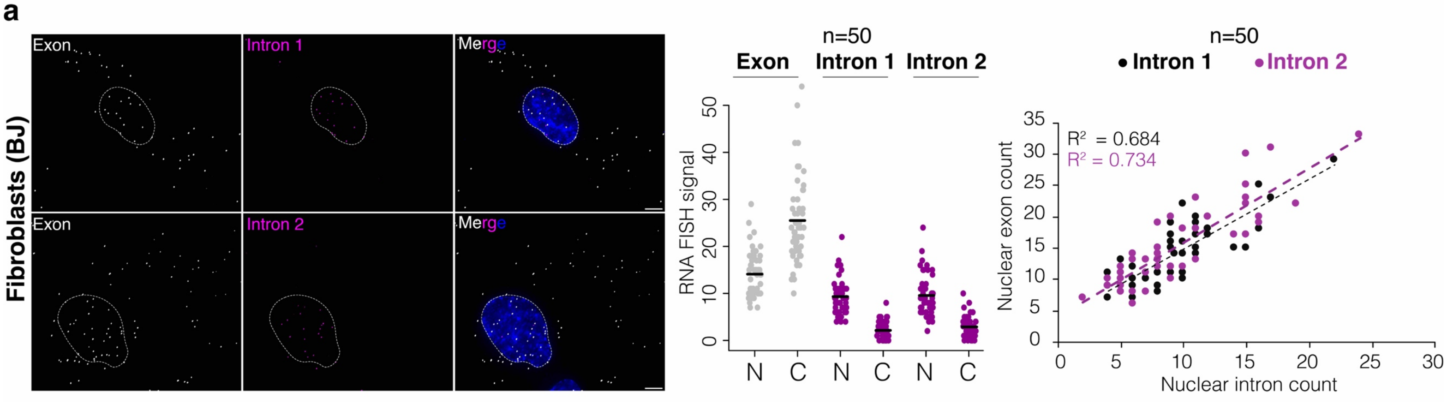


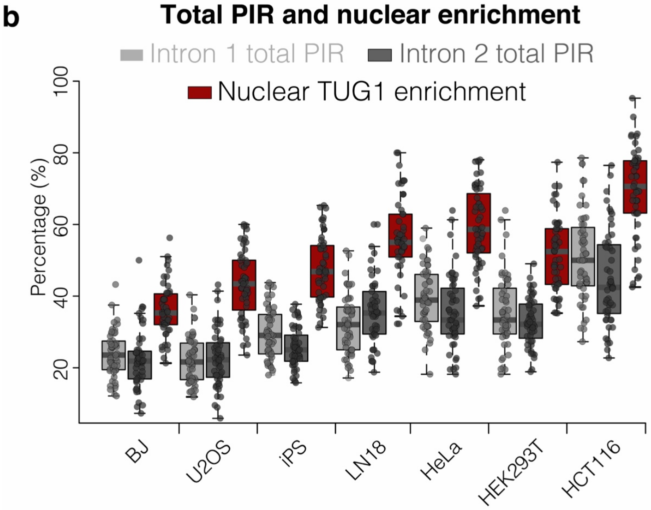


**Extended data figure 2: a**, Maximum intensity projections of representative images of TUG1 exon/intron smRNA FISH on human foreskin fibroblasts (BJ). Exon in gray, intron 1 and intron 2 in magenta. Nucleus in blue and outlined with a dashed line. Scale bar, 5 µm. Middle: quantification (*n* = 50) of spliced and unspliced transcripts for each intron in the nucleus (N) and cytoplasm (C). On the right: correlation between nuclear intron count and quantity of nuclear TUG1; intron 1 in black, intron 2 in magenta. **b**, Total percentage of intron retention (total PIR) of each intron and percentage of nuclear enrichment of TUG1 (nuclear TUG1 over total cell TUG1) across different cell lines.

*Extended Data Figures*

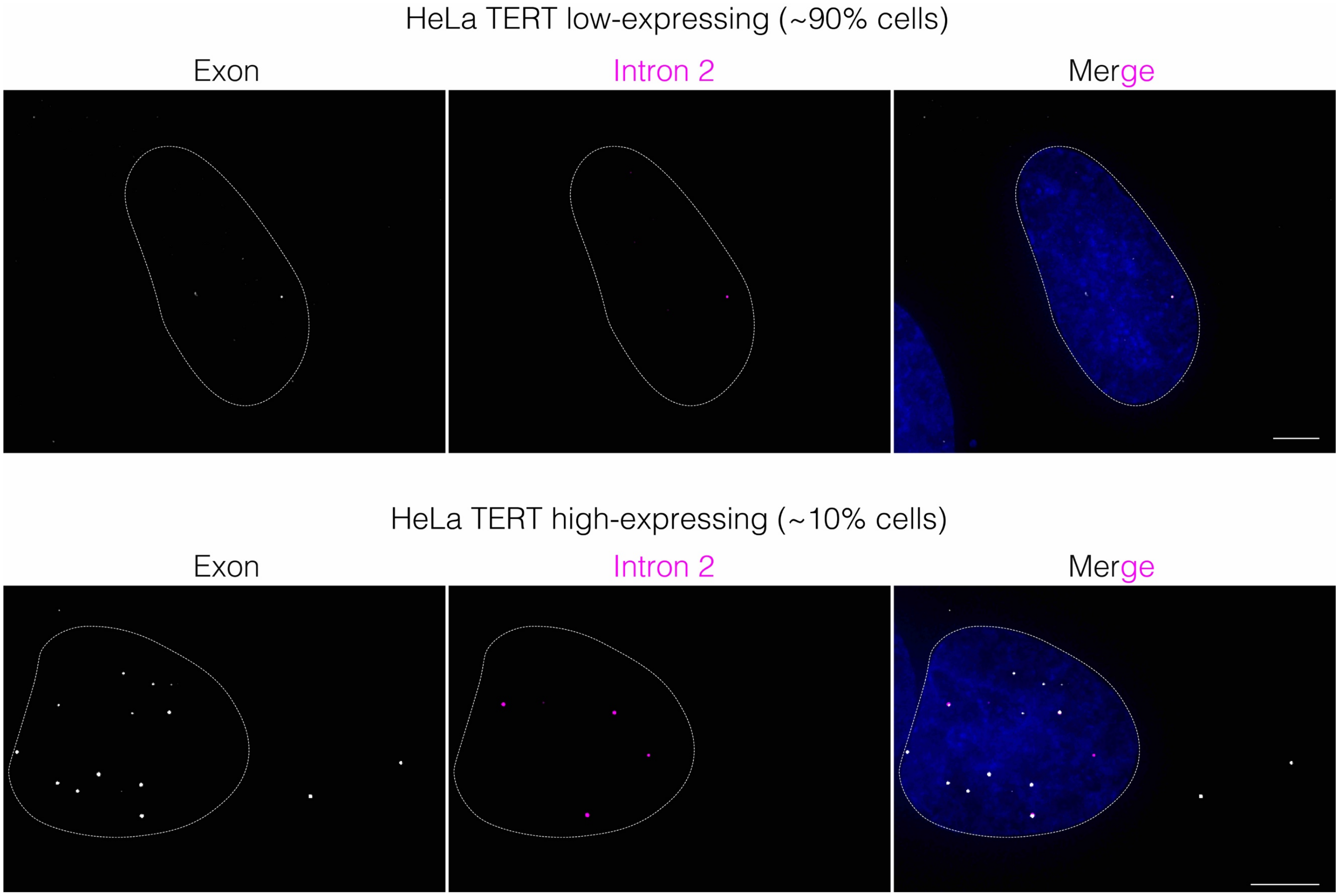


**Extended data figure 3:** Maximum intensity projections of representative images of TERT exon/intron 2 smRNA FISH on HeLa cells. Exon in gray, intron 2 in magenta. Nucleus in blue and outlined with a dashed line. Scale bar, 5 µm. On top shown TERT low-expressing cell, on bottom TERT high-expressing cell.

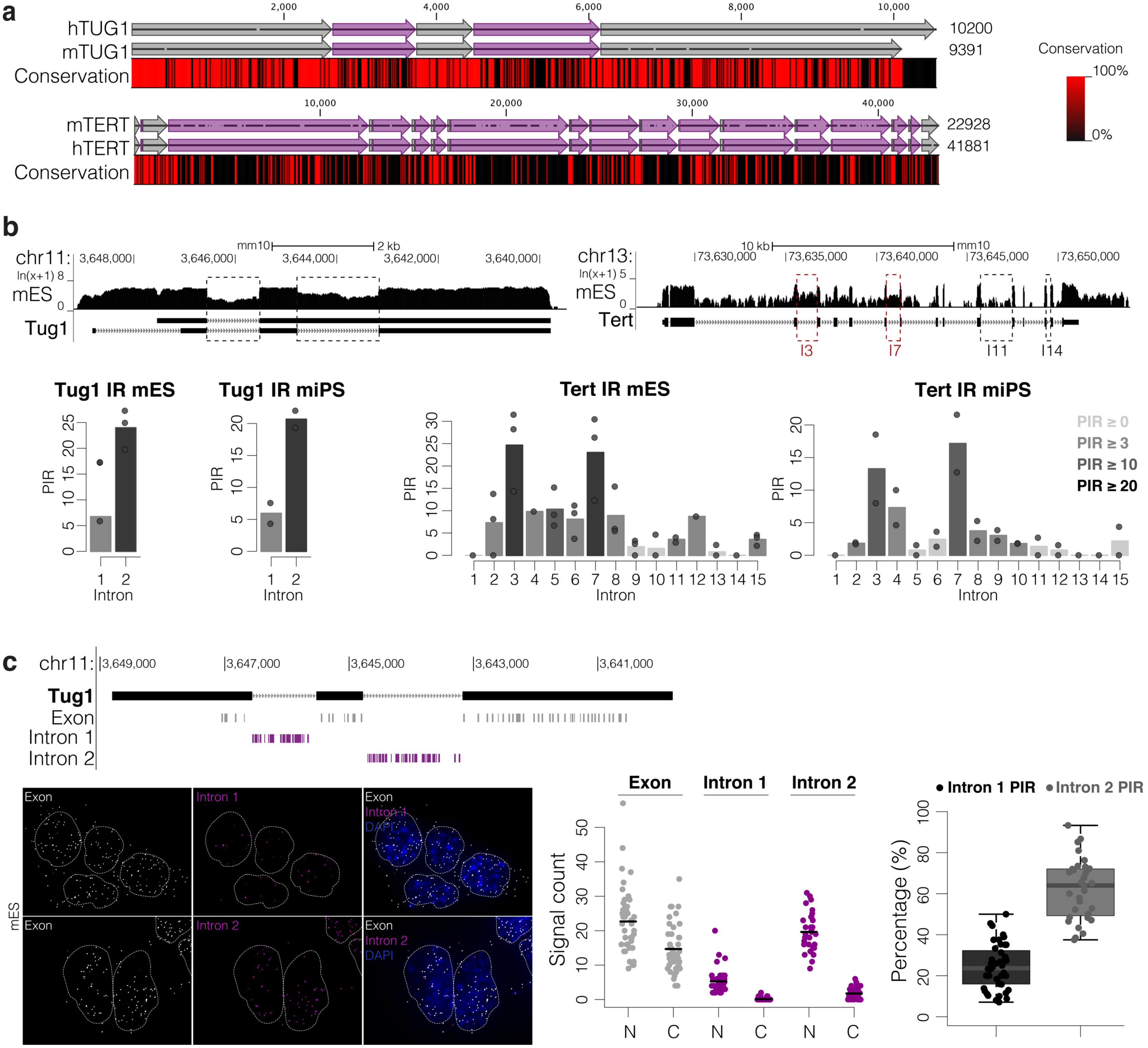


**Extended data figure 4: a**, Alignment of human and mouse TUG1 locus (on top) and TERT locus (on bottom). Exons depicted with gray arrows, introns with magenta arrows. Below the alignment is shown a conservation heatmap of conserved (red) and non-conserved (black) nucleotides. **b**, USCS Genome Browser showing RNA-seq coverage from mouse embryonic stem cells (mES) across the Tug1 locus and Tert locus. Below, percentage intron retention (PIR) of Tert (right) and Tug1 (left) in miPS and mES cells obtained with vast-tools analysis on RNA-seq data. Bars indicate means across replicates and dots individual replicates. **c**, Maximum intensity projections of representative images of Tug1 exon/intron smRNA FISH on mES cells. Exon in gray, intron 1 and intron 2 in magenta. Nucleus in blue and outlined with a dashed line. Scale bar, 5 µm. Middle: quantification (*n* = 30) of spliced and unspliced transcripts for each intron in the nucleus (N) and cytoplasm (C). On the right: percentage of nuclear intron retention (PIR) for intron 1 and intron 2.

*Extended Data Figures*

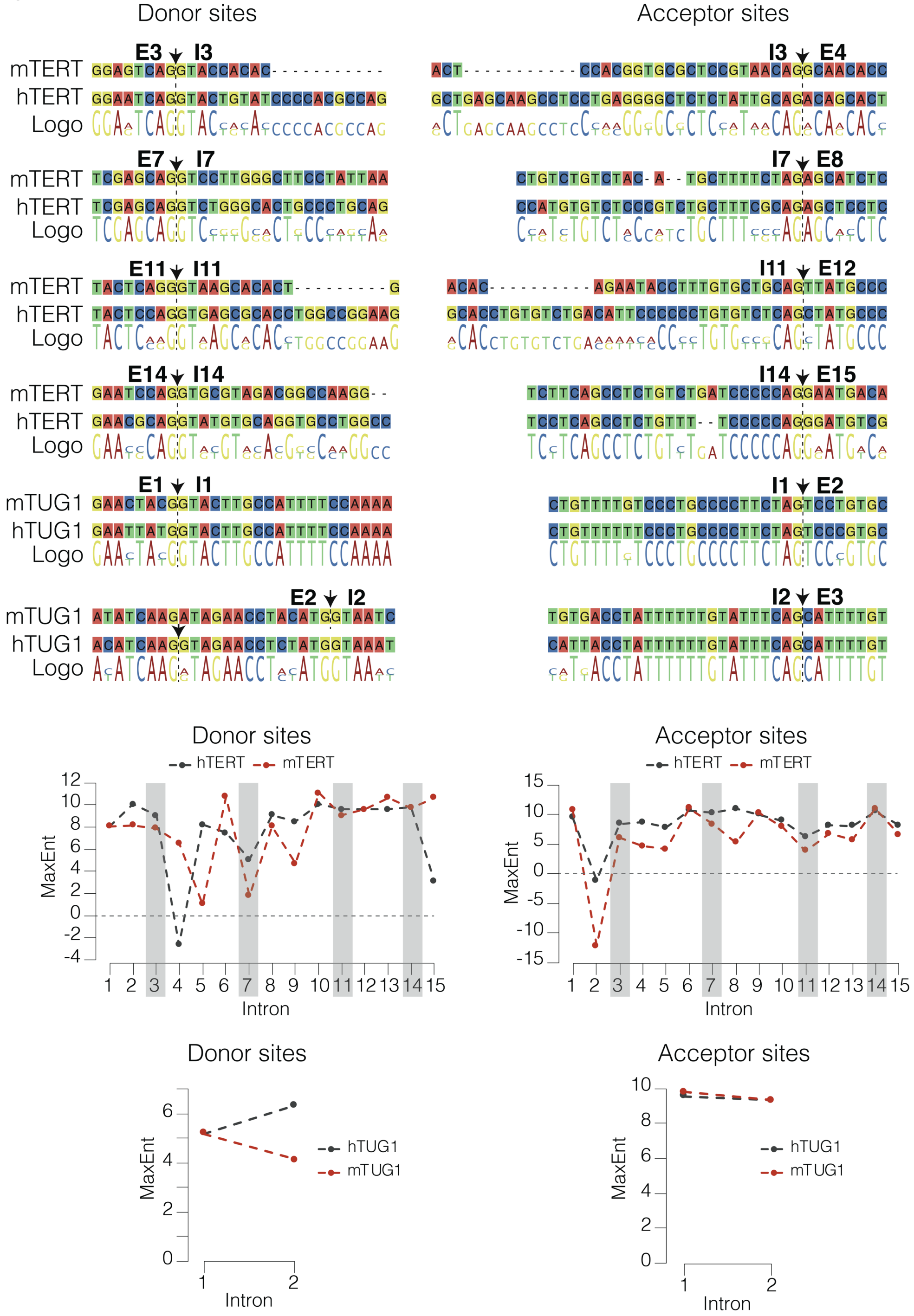


**Extended data figure 5:** Alignments of donor and acceptor splice sites and surrounding regions between human and mouse TUG1 and TERT. Arrows and dashed lines indicate the splice site. Mouse Tug1 E2/I2 junction is downstream of the E2/I2 junction in human TUG1. Below, comparison of donor and acceptor splice site strength measured using maximum entropy (MaxEnt) of human and mouse TUG1 and TERT retained and constitutively spliced introns.

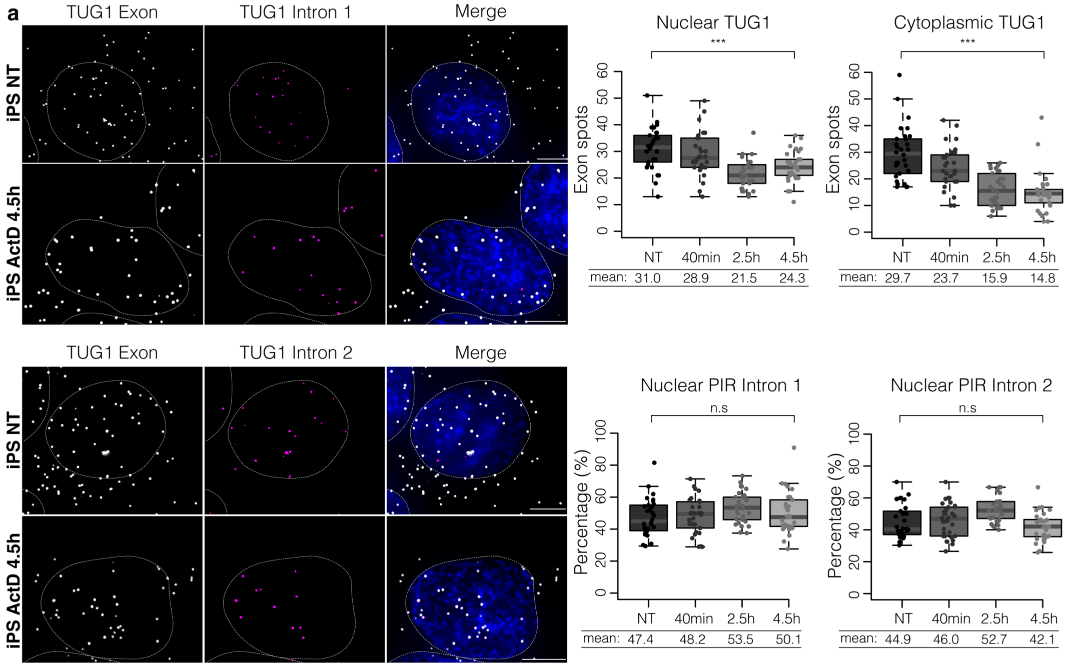


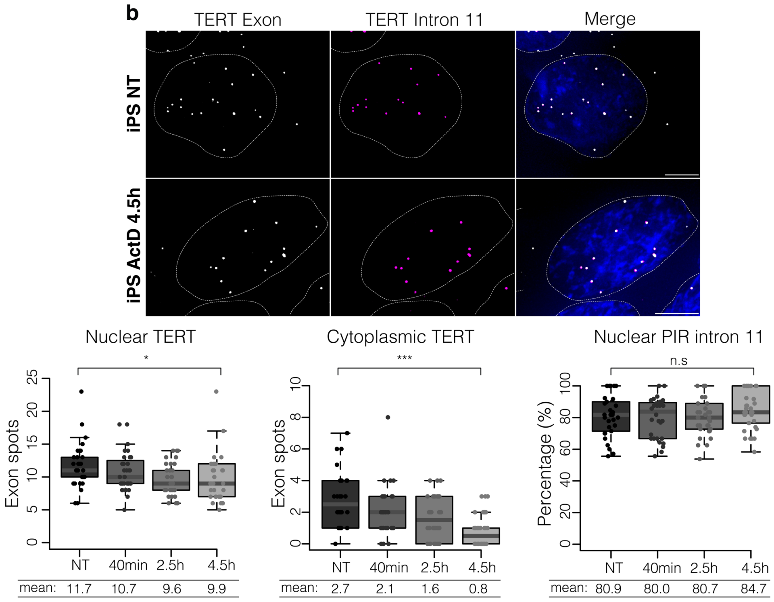


**Extended data figure 6: a**, Maximum intensity projection of smRNA FISH on iPS cells targeting TUG1 exon (gray) and intron 1 (magenta) or intron 2 (magenta) at time point 0 (NT) and 4.5 h after ActD treatment. Scale bar, 5 µm. Below quantification (*n* = 30-35) of smRNA FISH at each time point of spliced and unspliced TUG1 transcripts in the nucleus and cytoplasm; and percentage of nuclear intron retention (PIR) for intron 1 and intron 2 at each time point. n.s. = not significant, **P* ≤ 0.05, ****P* ≤ 0.001, as evaluated by unpaired *t*-test versus NT; *n* = 30-35 cells). **b**, Maximum intensity projection of smRNA FISH on iPS cells targeting TERT exon (gray) and intron 11 (magenta) at time point 0 (NT) and 4.5 h after ActD treatment. Scale bar, 5 µm. Below quantification (*n* = 30-35) of smRNA FISH at each time point of spliced and unspliced TERT transcripts in the nucleus and cytoplasm; and percentage of nuclear intron retention (PIR) for intron 11 at each time point. n.s. = not significant, **P* ≤ 0.05, ****P* ≤ 0.001, as evaluated by unpaired *t*-test versus NT; *n* = 30-35 cells).

*Extended Data Figures*

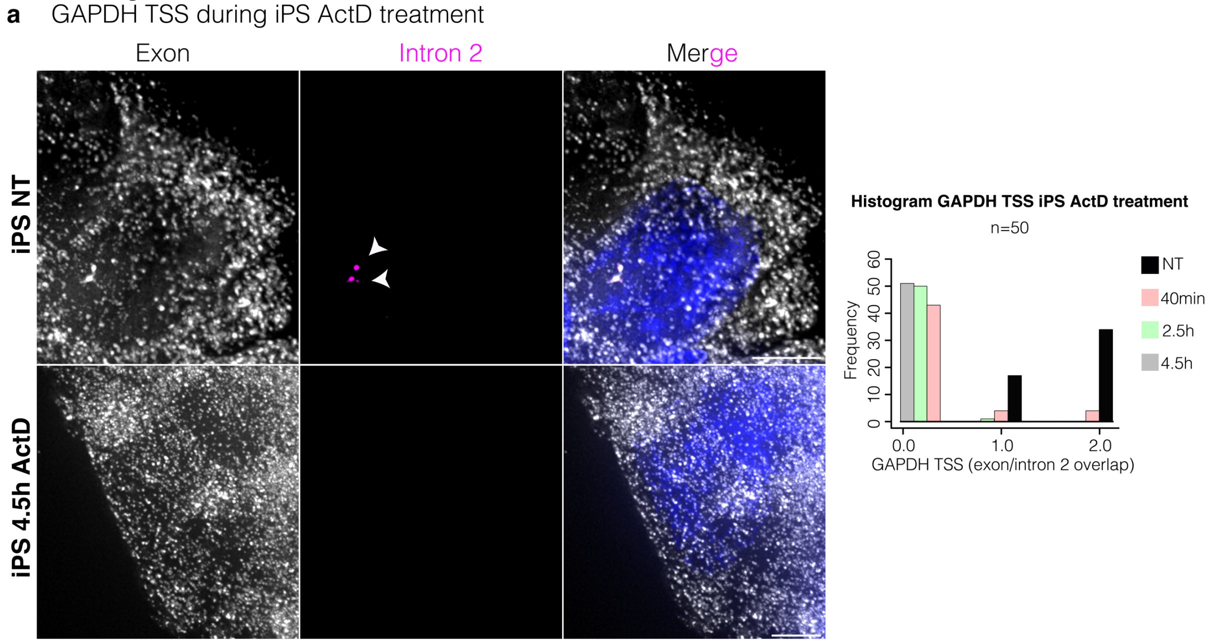


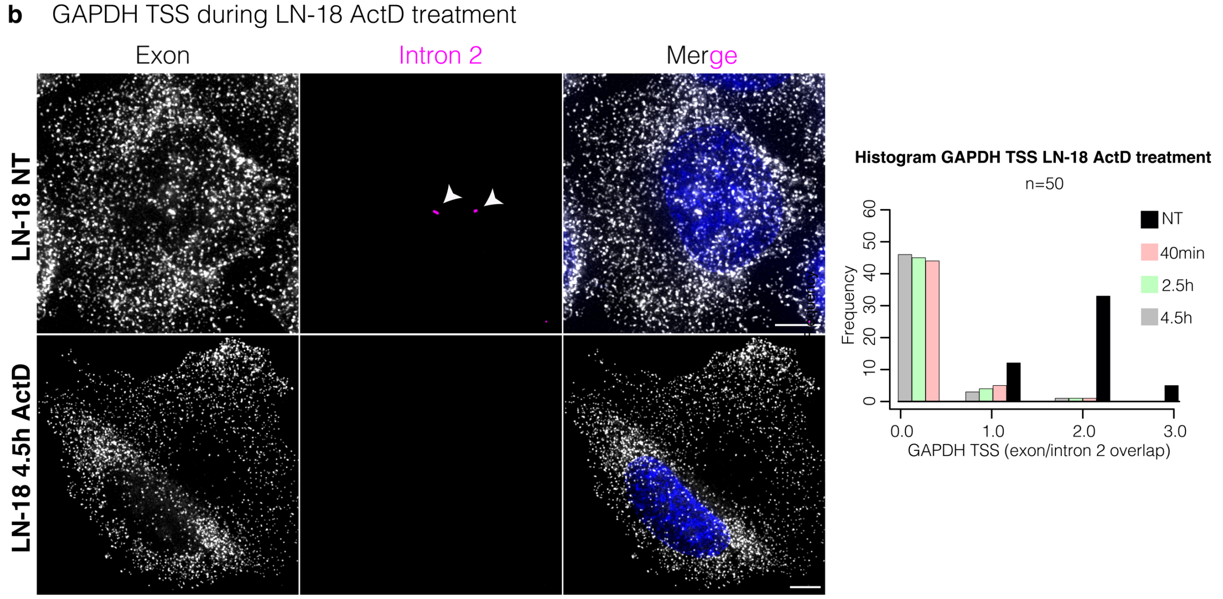


**Extended data figure 7: a**, GAPDH active transcription sites (exon/intron2 overlap) during ActD treatment of iPS cells. On the left, maximum intensity projection of GAPDH exon (gray) and intron 2 (magenta) smRNA FISH on untreated (NT) and 4.5 h of ActD treatment iPS cells. Scale bar, 5 µm. On the right, histogram showing the distribution of GAPDH transcription sites during the ActD time course treatment. **b**, GAPDH active transcription sites (exon/intron2 overlap) during ActD treatment of LN-18 cells. On the left, maximum intensity projection of GAPDH exon (gray) and intron 2 (magenta) smRNA FISH on untreated (NT) and 4.5 h of ActD treatment LN-18 cells. Nucleus in blue. Scale bar, 5 µm. On the right, histogram showing the distribution of GAPDH transcription sited during the ActD time course treatment.

*Extended Data Figures*

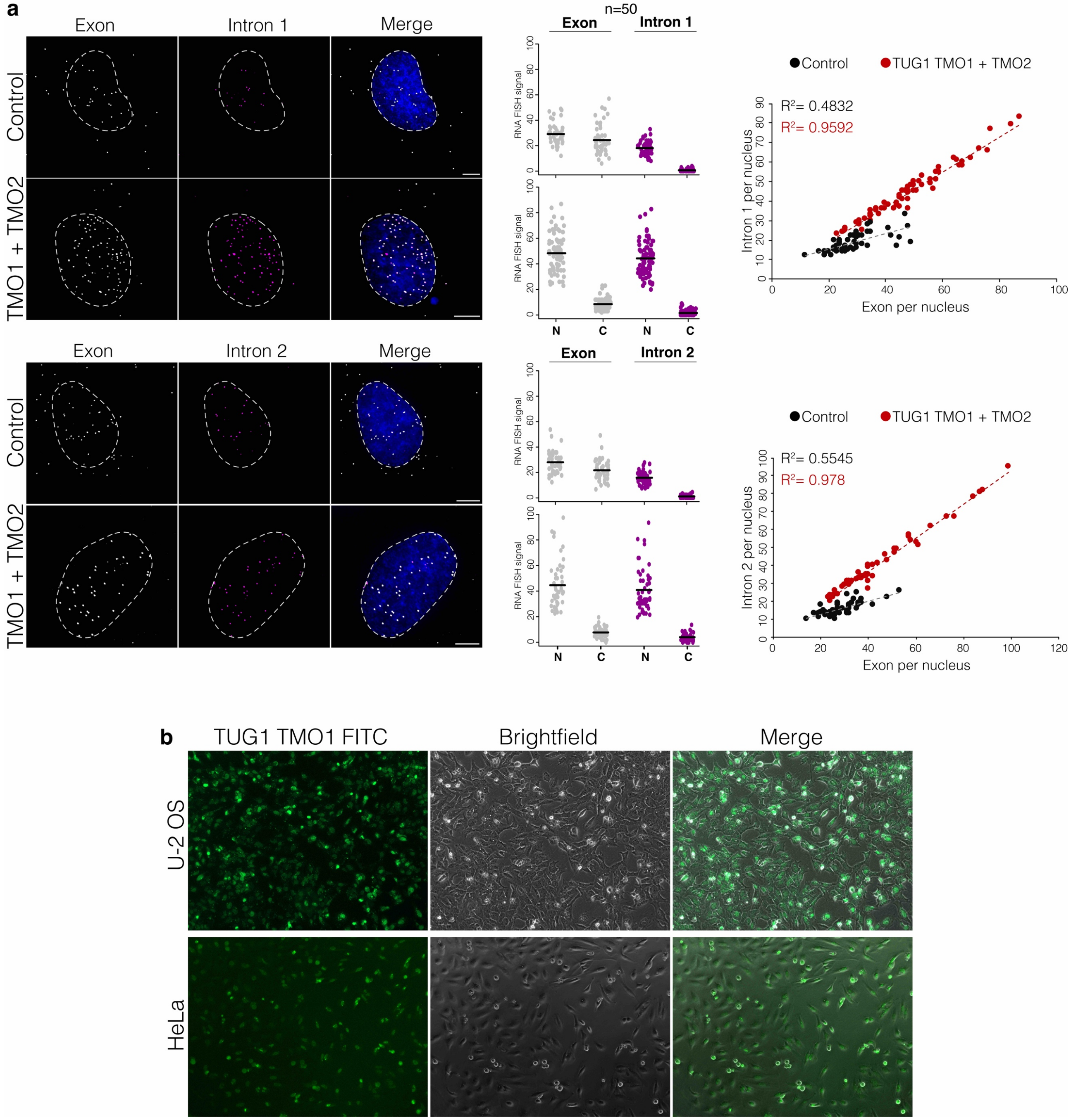


**Extended data figure 8: a**, Maximum intensity projections of TUG1 exon (gray) and intron 1 (magenta) or intron 2 (magenta) smRNA FISH on HeLa cells transfected with control TMO and with TUG1 TMO1 and TMO2. Nucleus in blue, outlined with a dashed circle. Scale bar, 5 µm. Towards the right, quantification of spliced and intron-retained TUG1 for each intron in the nucleus (N) and cytoplasm (C). Shift in the quantity of nuclear TUG1 that retains introns shown on the right, TUG1 TMO1 and TMO2 in red, control TMO in black (*n* = 50-80 cells). **b**, U-2 OS and HeLa transfection efficiency assessed by TUG1 TMO1 labeled with FITC intake (in green).

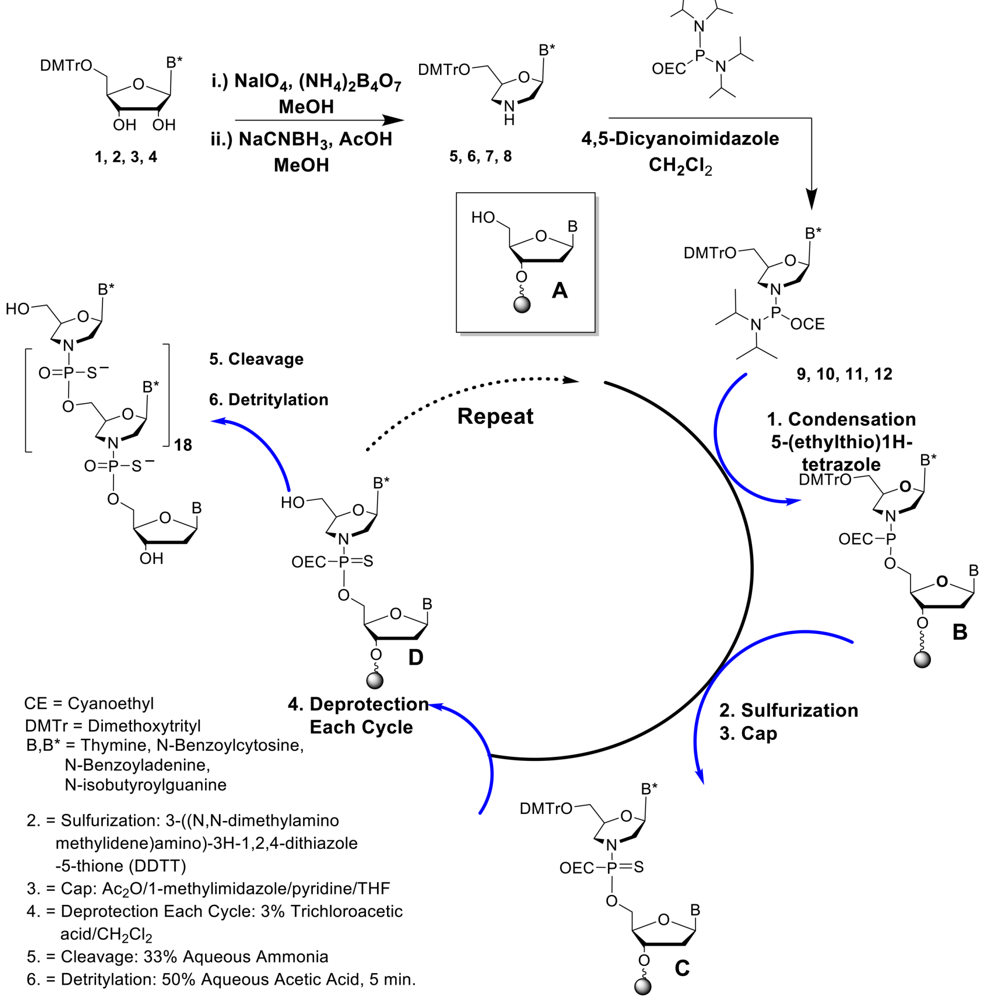


**Extended data figure 9: Synthesis of thiomorpholino oligonucleotides**. Synthesis of thiomorpholino oligonucleotides. Morpholino nucleosides 5-8 and morpholino phosphorodiamidites 9-12 were synthesized starting from appropriately protected ribonucleosides 1-4 as shown in the top part of the figure. The synthesis cycle begins with detritylation of succinyl CPG 500 supported nucleoside (A). Condensation with phosphorodiamidites 9-12 generates B, which is sulfurized to produce the thiophosphoramidate morpholino triester (C). After capping the failures, detritylation is carried out to produce (D), which is then ready for the next synthesis cycle. Cleavage of final TMO oligonucleotide was carried out using 28% aqueous ammonia at 55°C for 16 h.

*Extended Data Figures*

**Extended data Table 1:** TERT and TUG1 PIR of each intron in hiPS, mES and miPS cells calculated with Vast-tools on RNA-Seq.

**Extended data Table 2:** Length and GC content of human and mouse retained TERT and TUG1 introns.

**Extended data Table 3:** Global PIR analysis for coding and lncRNA genes in hiPS cells calculated with Vast-tools on RNA-Seq.

**Extended data Table 4:** Global PIR analysis for coding and lncRNA genes in mES and miPS cells calculated with Vast-tools on RNA-Seq.
